# Mapping Structural Drivers of Insulin Analogs Using Molecular Dynamics and Free Energy Calculations at Insulin Receptor

**DOI:** 10.1101/2022.05.27.493461

**Authors:** Mohan Maruthi Sena, C Ramakrishnan, M. Michael Gromiha, Monalisa Chatterji, Anand Khedkar, Anirudh Ranganathan

## Abstract

A century on from the discovery of insulin, a complete understanding of insulin interactions with the insulin receptor (IR) at atomic level remains elusive. In this work, we have leveraged recent advancements in structural biology that have resulted in multiple high-resolution structures of the insulin-IR complex. As a first step, we employed molecular dynamics (MD) simulations to unravel atomic insights into the interactions between insulin-IR complexes in order to better understand ligand recognition at the receptor. The MD simulations were followed up with free energy perturbation (FEP) calculations to discriminate between and elucidate the drivers for ligand association for various natural and man-made insulin analogs. As an example, these calculations were utilized to understand the molecular mechanisms that characterized the loss-of-function seen in disease-associated insulin mutations seen in different populations. Further, multiple man-made insulin analogs spanning a range of potencies, mutations, and sequence lengths were studied using FEP and a comprehensive molecular level map of potency determinants were established. ∼85% of FEP calculations captured the direction of shift of potency, and in ∼53% of cases the predictions were within 1 kcal/mol of experiment. The impressive accuracy of FEP in recapitulating functional profiles across such a span of insulin analogs and potency profiles provided clear evidence of its utility in computational mutagenesis. In addition to the impressive accuracy, the ability of FEP to provide a dissected understanding of protein residue, solvent and solvent-mediated contributions to binding energy clearly establishes its utility in the design of novel insulins and peptides in general.

## Introduction

Insulin is an anabolic peptide hormone produced by β-cells of the islets of Langerhans within the pancreas that plays a crucial role in glucose homeostasis.^1–5^ Mature insulin is stored in the hexameric form within the pancreas^6–8^ and is produced via enzymatic cleavage of proinsulin, which contains a C-chain peptide connecting A and B-chains.^9–11^ The C-peptide is cleaved by the specific endopeptidases resulting in the mature form of the insulin.^12,13^ Activity in vivo involves dissociation of the storage hexameric form initially into dimers and then to a monomer which engages with insulin receptor (IR).^9,14^ Monomeric bioactive human insulin has been structurally well-characterized and is a 51 amino acid peptide consisting of two chains A and B with 21, 30 amino acids respectively.^9,15,16^ The chains are held together by two inter-chain disulfide bridges(A7-B7; A20-B19) and further stabilised by an intra-A-chain cysteine linkage (A6-A11).^15,17,18^ Abnormalities in insulin secretion or resistance to insulin leads to type-1 or 2 diabetes.^15,18–21^

Monomeric insulin mediates a host of important functions primarily through its engagement with the IR.^1,18,22,23^ The IR belongs to the receptor tyrosine kinase (RTK) family of proteins that include type 1 and 2 insulin like growth factor receptors (IGF1- and IGF-2R) and the orphan insulin receptor-related receptor.^15,18,22^ The RTKs share a common architecture consisting of an extracellular domain (ECD), a transmembrane segment and an intracellular kinase domain.^15,18,20^ The RTKs form homodimers as quaternary structures and undergo conformational rearrangements upon ligand binding that results in auto-phosphorylation of specific tyrosine residues intracellularly and downstream signaling.^18,23^ Key physiological functions mediated by IR activation include glucose uptake, glycogen synthesis in skeletal muscles, diminished glucose synthesis in hepatocytes and lipid degradation in adipocytes.^1,15,18,19,21^ These roles of insulin-mediated IR activation in metabolism and homeostasis make them primary targets for metabolic disorders such as diabetes that affect millions of lives globally.^19,21,22^

Insulin has been well-studied for nearly a century but a molecular level understanding of its receptor binding and subsequent signaling remains unclear.^18,21,24^ For a long time this was due to lack of high resolution structures of insulin bound to the IR. A key structural breakthrough was achieved when McKern *et al*., provided us with the first glimpse of the IR ectodomain and revealed an inverted ‘V’ like conformation in the absence of insulin.^23^ In 2013, the first high resolution structure of an insulin-IR complex revealed a single insulin molecule bound to the extracellular domain of the IR.^18^ This structure showed that the high-affinity (HA) binding site of insulin lies between the leucine-rich L1 domain and the C-terminus of the α-domain (α-CT).^18^ In 2018, single electron microscopy (EM) experiments have shown that activation of the insulin-IR complex involved transitioning from an inverted ‘V’ conformation to a T-like state that is thought to allow the transmembrane domains to come in proximity to each other and result in auto-phosphorylation and downstream signaling.^25^ During the course of this work, other structures have demonstrated the presence of the hypothesised low-affinity insulin binding sites at the IR which have revealed the complexities of ligand-receptor engagement (Table S1). _2,2,18,26,26–29_

In contrast to IR-insulin complexes, the structure of free insulin has been known for decades.^6,7,30–32^ Multiple computational studies have focused on understanding the conformational dynamics of the molecule in the monomeric, dimeric and Zn^2+^ mediated hexameric states.^33–39^ These include the use of computational alanine scanning, free energy calculations with MMGBSA, study of solvent mediated effects, and enhanced sampling methods to understand the conformational spread within the insulin molecule.^39–42^ In parallel, Kiselyov *et. al* formulated a harmonic oscillator model to explain empirical results from insulin binding experiments with IR and IGF-1R.^43^ Previous in silico studies of insulin behavior in complex with IR relied on docking generated models of the bound state.^33^ However, experimental structures with insulin bound revealed differences between the docking solutions used in those studies, with the actual bound conformation of insulin.^18,25–29,44^

Whereas, the previous computational studies provided important glimpses into the function of insulin at a molecular level, a complete picture of insulin activity at the IR remains elusive. We leveraged new structural information that has emerged in the last few years with regards to insulin-IR complexes to build a deeper understanding of the hormone function (Table S1).^2,2,18,28,29,45^ In this study, we used MD simulations along with an adaptation of the powerful free energy perturbation (FEP) method to perform in silico mutagenesis, in order to map and understand the contribution of different residues and domains on receptor binding and activation.^46^ FEP has been widely used to compute relative protein-ligand binding affinities for small-molecules and mapping the effect of mutating receptor residues,^46^ however, to the best of our knowledge this is the first use of the method for assessing mutations in bound polypeptide ligands, such as insulin.^47–49^ Unlike, with other faster free energy estimation methods, the use of FEP allows for deconvolution of the energy contributions of individual residues and can shed light on the role of waters due to the use of explicit solvation. Hence, in this work, MD and FEP are used as the foundations to understand the structural drivers of insulin activity and function via the IR.

In order to probe the utility of MD and FEP in enhancing our understanding of insulin engagement with its receptor we divided our study into three defined segments. Initially, we provide a baseline for ligand stability in the binding site and map out key interactions between WT-Ins and IR using short MD simulations. Next, we looked at naturally occurring disease-linked variants of human insulin to see if we can understand the molecular drivers behind the loss-of-function seen with these mutants using FEP. Finally, we extended the FEP calculations to study a variety of well-characterized man-made insulin variants in order to link amino acid residue changes to the different potency profiles that they display. These calculations help us to provide the relationship between observed metabolic potency and correlate it with the corresponding atomic changes. This atomic level information could be used to designing new insulin variants. In addition, a recapitulation of functional profiles across different analog types would act as a demonstration of the generalizability of the FEP method as a tool for performing in silico mutagenesis towards the design of novel peptides and proteins. In this article, we use the term “ligand” as nomenclature in place of “insulin” or “insulin analog.”

## Results & Discussions

### Analysis of insulin bound IR structures

IR is a functional homodimer consisting two α-chains and two β-chains that form heterodimers represented as (αβ)_2_ that contains extra-cellular, transmembrane and kinase domains.^5,15,18^ To date, EM and XRD have both provided high-resolution structures of both IR in complex with their cognate peptide hormones, which broadly capture the homodimers in variations of the inverted ‘V’ or “T” conformations (Table S1).^18,26,28,29,45^ Recently IR cryo-EM structure of insulin bound to IR ectodomain (ECD) elucidated the mechanism of the activation upon ligand binding.^50,51^ Upon binding of the insulin ligand to the receptor, a conformational transformation occurs from inverted ‘V’ to ‘T’ shape which brings the L1, α-CT domain of one dimer (αβ), and FnIII-1 loop of another dimer (αꞋβꞋ) other close to each other.^45^ For this study, we chose the EM derived IR-insulin complex (PDB accession code: 6CE9;) as our reference structure.^45^

In IR, the wild type insulin (WT-Ins) symmetrically can bind to four binding sites (site 1, 1’, 2 and 2’) spanning the active dimeric insulin receptor.^2,26,27^ Two of them (1 and 1’) are designated as high affinity (HA) sites with the other two (2 and 2’) labelled as low affinity (LA) sites. The labelling of the insulin binding sites in the IR as HA and LA sites stems from the large differences in insulin surface burial upon binding to these distinct sites (706 Å^2^ in HA site and 394 Å^2^ in LA site, Table: S2)^26^, which in turn leads to differences in binding affinities. The L1 of one subunit and α-CT’ and FnIII-1’ of another subunit form the HA binding site (Figure 1).^18,25,26,28^. It is to be noted that in this study, we have focused on the receptor high affinity (HA) sites. From the experimental structures available for human insulin in complex with IR, we observed that the insulin mainly adopts an “open” conformation in the HA sites when the overall receptor is in the “T” conformation (active) (Figure 1). The important structural features associated with the open conformation are a different conformation of the cysteines that form the two inter-chain disulfide bridges and the unwound conformation of C-termini of B-chain when compared to the unbound insulin structure.

**Figure 1.**
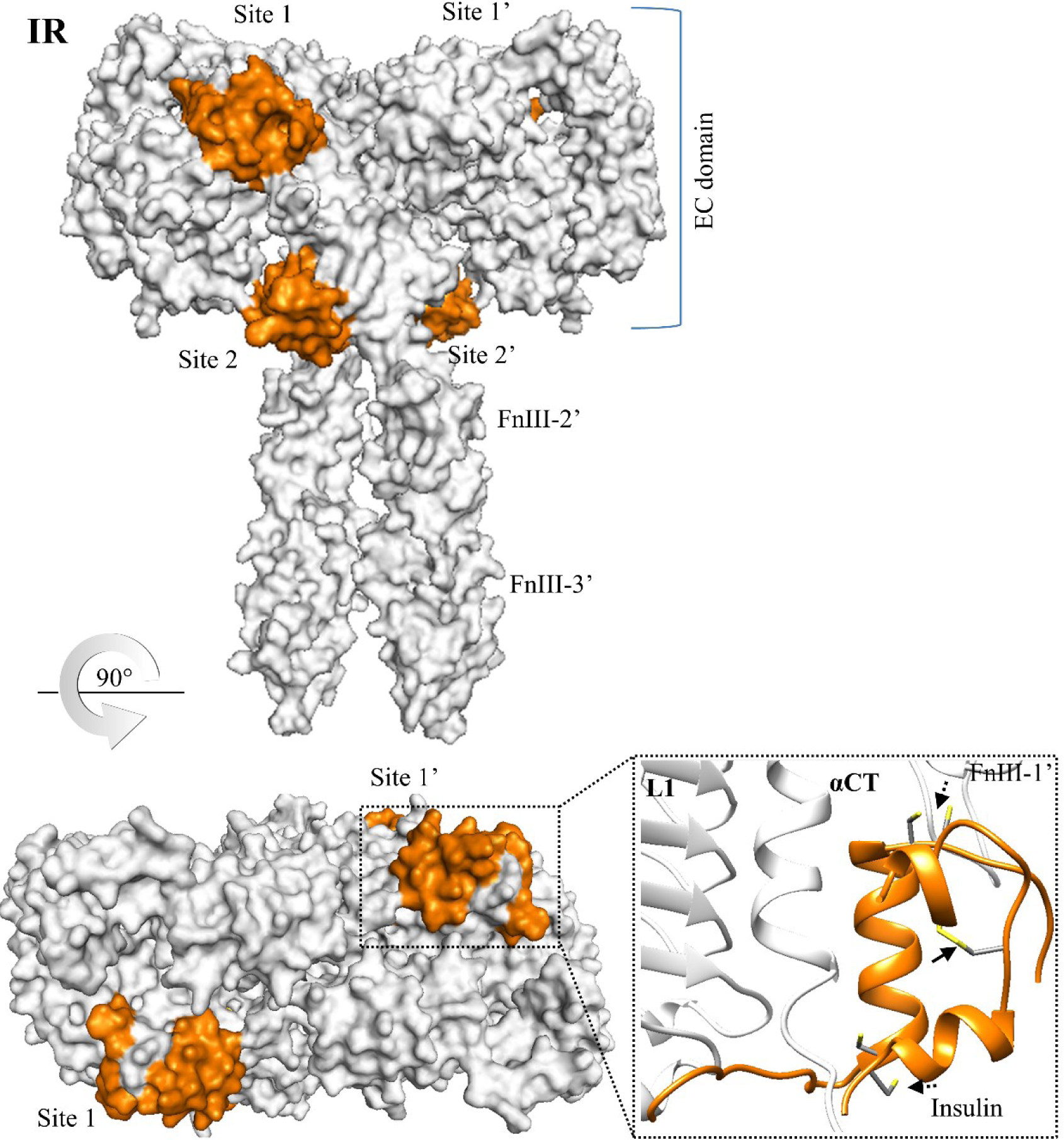
Structural analysis of experimentally obtained structure of IR: Top panel - The experimentally obtained structure of the IR receptors used in this work are shown in the surface representations with their respective ’T’ conformations (active). The binding sites for the receptor are highlighted in orange. For IR, the high and low affinity sites are labeled as 1/1’ and 2/2’, respectively. Bottom panel – A zoom in of the high-affinity sites of the receptor are shown with the insulin molecule bound (represented as an orange cartoon). Arrows depict the intact inter- and intra-chain disulfide bonds in the complex.

### Short MD simulations of the IR with WT-Ins to assess stability and ligand engagement

To assess overall stability of the complex and peptide engagement with the HA sites of IR, simulations of the IR-WT-Ins complex were performed using all atom MD of a fully hydrated system at 300 K (Figure S1a). The starting coordinates for the simulations were obtained from the active state of insulin-IR complexes from Scalpin *et al.,* [PDB accession code: 6CE9].^45^ Missing segments of the receptor structures were modelled using homology modelling.^52^ After initial restrained equilibration, the entire system was simulated with complete flexibility.

The interaction framework provided by the initial structures was overall well-maintained throughout the trajectory and the system was largely stable (Figure 2a: RMSD). The low average fluctuation for each peptide ligand further elucidated the intrinsic stability of the bound conformation. In addition, the solvent accessible surface area (SASA) was also calculated for each ligand and the obtained averages were in line with those reported for the HA site of the EM structures, further reiterating the stability of the complexes, (Table: S2).^45^

**Figure 2:**
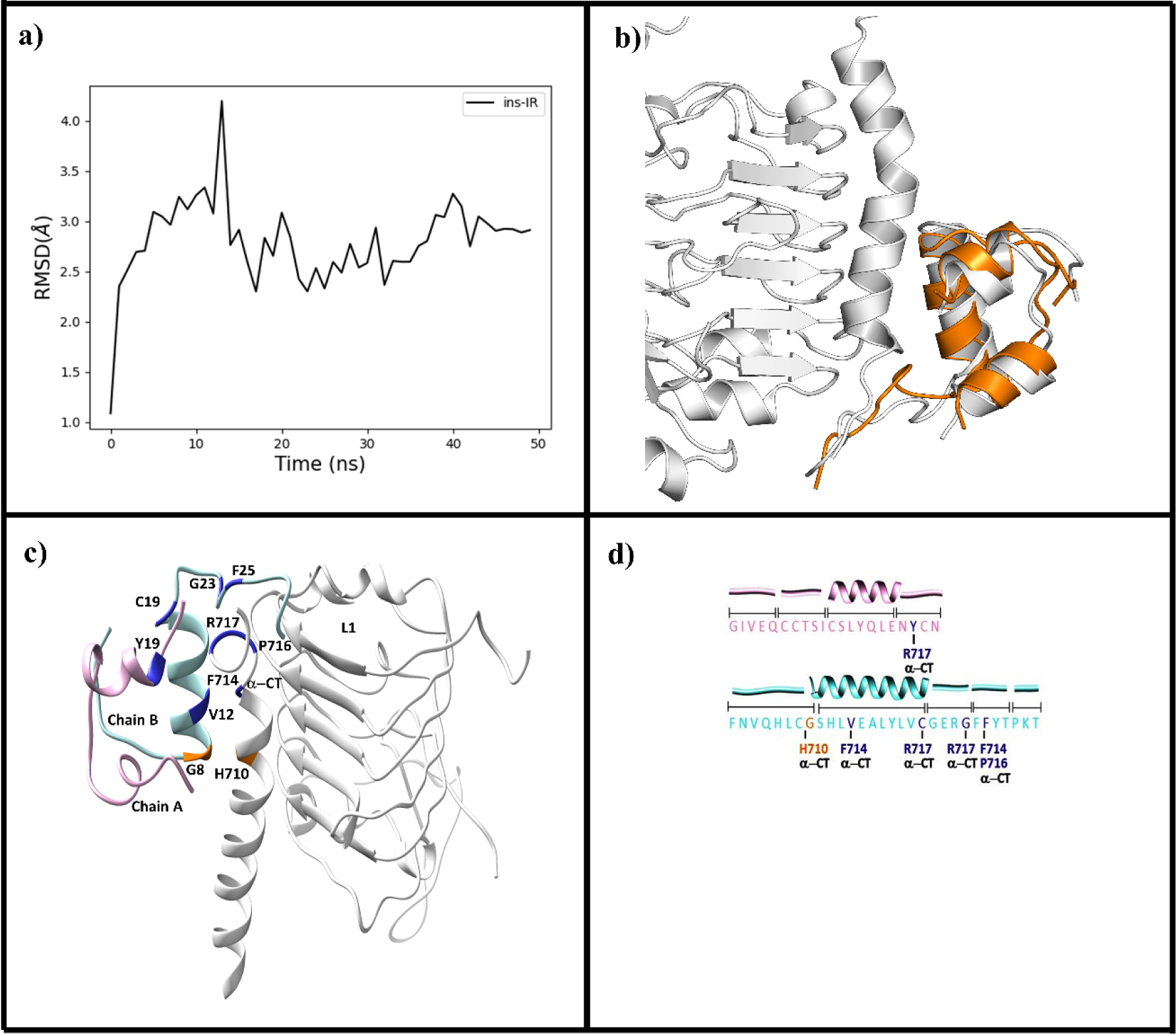
Assessment of stability of insulin-IR complex from MD simulations with the starting coordinates as reference a) Cα-RMSD from the simulation with respect to the initial structure, insulin-IR (black), b) insulin-cognate complex: the insulin starting structure (grey cartoon) is shown in alignment with the average structure of insulin (orange cartoon) and receptor (grey cartoon) obtained from MD simulations. c) Analysis of stable contacts of WT-Ins in IR receptor with more than 90% lifetime obtained from MD simulations. Ligand-receptor contacts were defined using a distance cutoff of 4.5 Å from any ligand residue. The respective receptors in complex represented as grey cartoons, the chain-A of the WT-Ins is color coded in Mauve color, whereas chain-B is color coded in cyan. The interacting residues from ligand and receptor are identically color-coded domain-wise and domain names/residue names are further highlighted in text. d) Full insulin sequence is shown in the text and the corresponding interactions/domains from IR were highlighted using text.

To better understand the ligand-receptor interaction networks, we analysed intermolecular interactions within 4.5 Å distance cut-off around the insulin. When an occupancy cut-off of 90% was applied to the interactions within a 4.5 Å shell around the ligand, the analysis revealed that highly stable interactions are dominated by those between the C-termini of both A- and B-chains of the respective ligands with the α-CT domain of the receptor, (Figure 2c). Insulin engages IR-α-CT extensively with its B-chain. In the highly conserved aromatic B-chain or equivalent domain of both ligands, Insulin engages primarily with B24Phe, (Figure 2c). Whereas, interactions with the α-CT were still dominant, engagement with the Fn-III and L1 domains were also established. (Figure 2c). In the Chain-A of insulin, the short loop segment with residues A6-A10 is found to interact with Fn-III-1’ domain of the receptor whereas helical segment A11-A17 does not form long-lived interactions (>90% residence time) with the receptor. In the B-chain, the helical stretch between the residues B8-B20 interacts with L1-domain of the receptor in addition to α-CT and Fn-III segment, (Figure: 2d). The short hinge region (GXXG) interacts with the R717 of the α-CT domain. Overall, the analysis reiterated the importance of the insulin B-chain and C-terminus of the A-chain along with the α-CT domain of the IR in anchoring the intermolecular interactions in the HA binding site.

### Understanding the loss-of-function associated with disease causing mutations

Three missense variants in insulin have been reported to cause hyperinsulinemia or hyperproinsulinemia,^53–55^ and carriers were observed to develop insulin resistance that could lead to a diabetic state.^53^ Three natural mutations that have been observed in different populations, namely insulin Wakyama (ValA3Leu), insulin Chicago (PheB25Leu), and insulin Los Angeles (PheB24Ser). These mutations confer reduced biological activity towards IR with Wakayama displaying the largest loss-of-function.^53,56^

We aimed to use FEP to probe the molecular drivers responsible for the losses-of-function. For consistency, FEP calculations were performed on the HA sites of the IR using the same starting coordinates as our MD simulations [PDB accession code: 6CE9].^45^ These calculations were performed with spherical boundary conditions (Figure: S1), which overall maintained the receptors in the ligand-bound states represented by the starting structures (Figure 1). All atoms within the sphere were fully flexible whereas residues outside the sphere were tightly restrained to their initial coordinates, thereby maintaining the overall conformation of the receptor (Figure S1a). FEP schemes were devised for each disease mutant assessed in this study utilizing the fact that free energy is a state function and independent of path (Table S3). The number of intermediate steps for each transformation was assigned in order to minimize hysteresis. In total 1.8 μs of simulations was performed for these three mutants, with an average of ∼620 ns used for each FEP calculation for each receptor. Relative binding free energies (ΔΔG_bind_) were calculated for each mutant using a thermodynamic cycle with the free ligand in water acting as the reference state (Figure S1b). ΔΔG_bind_ with respect to WT-Ins or equivalent was used to quantify the relative affinities of each molecule to the IR in their respective conformation.

The ΔΔG_bind_ for insulin Wakayama from FEP calculations (+0.61 kcal/mol) captured the loss-of-function associated with this mutation albeit underestimated the degree of loss when compared to experiment (+3.1 kcal/mol) [Table S4, Figure 3].^53^ Decomposition of the ΔΔG_bind_ demonstrated that the annihilation of the larger alkyl chain of the Leucine residue (mutant) to valine (WT) restored favourable energies at the IR. Structural analysis further revealed that a water-mediated interaction network connecting D496 (Fn-III domain), K703 & D707(α-CT) is lost when the naturally occurring valine is changed to the bulkier leucine (Figure 3a). Similarly, the ΔΔG_bind_ for insulin Chicago was in good agreement with the loss-of-function associated with this mutant(+0.79 kcal/mol) with a similar underestimation when compared to experiment (+2.3 kcal/mol) [Table S4].^53^ Decomposition of ΔΔG_bind_ reveal the loss of affinity associated with the annihilation of phenyl ring atoms (WT) is not compensated by the gain from improved electrostatics in the mutant. Further structural insights revealed the loss of function for the insulin Chicago is due to subtle perturbations of the interaction network around B25Phe, when the aromatic Phenylalanine residue is replaced with a Leucine (Figure 4b).

**Figure 3:**
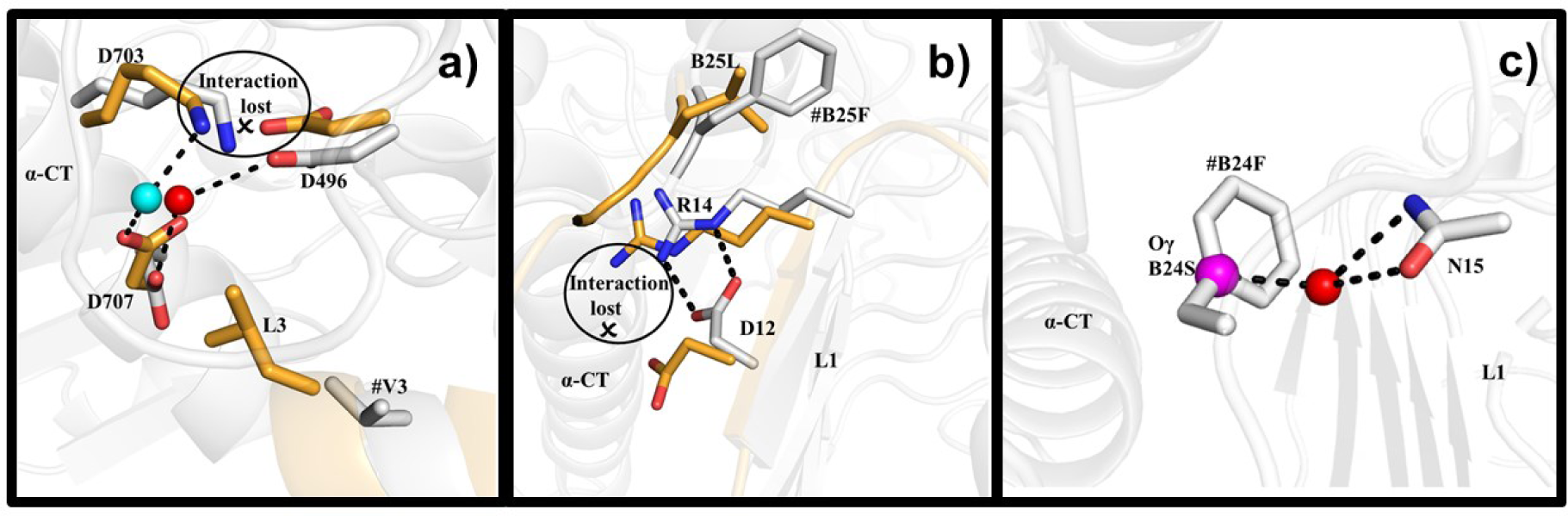
Structural changes accompanying disease-causing mutations are shown. Hydrogen bonds are indicated by dashed lines. In all figure panels WT-Ins-IR complex is shown with carbons in grey and the corresponding disease-mutant-IR complex is depicted with carbons in orange. In general the complexes are represented as cartoons with key residues from the receptors and ligands highlighted as sticks. **a**) insulin Wakayama, showing the loss of a water-mediated interaction network between two domains in the IR. The red sphere represents the bridging water in the WT-Ins-IR complex between 496Asp, 707Asp and K703Lys. The cyan sphere shows the displacement of this bridging water due to the larger Leucine residue at A3 position, which causes the loss of the inter-domain interaction with 496Asp in this complex; b) insulin Chicago the salt-bridge between 12Asp and 14Arg within the L1 domain is present in the WT-Ins-IR complex and is lost upon mutating B25Tyr to Leucine, and c) Insulin Los Angeles depicts the presence of B24Phe in a buried pocket within the IR. Mutation to the smaller serine resulted in partial solvation of this pocket, where the residue was observed to forms a water-mediated link to the 15Asn residue within the L1 domain.

**Figure 4:**
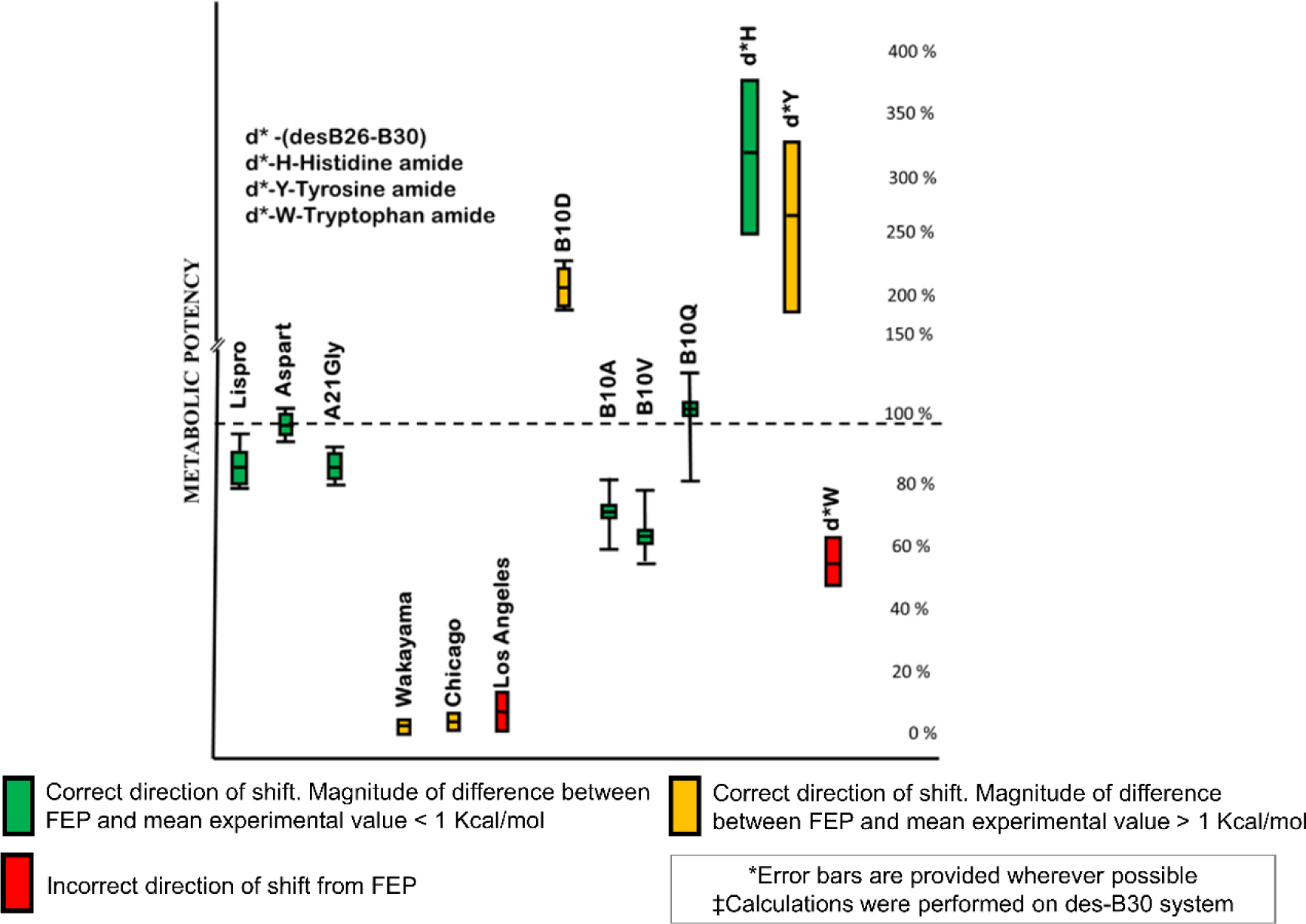
Summarizing the correspondence obtained between calculated values from FEP calculations & the relevant metabolic potencies experimentally values obtained from various studies for all the insulin variants evaluated in this study. The relative binding affinities ΔΔG_bind_ obtained from the FEP calculations at IR were compared with experimental metabolic potencies (left). The metabolic potencies of mutations are reported to relative to WT-Ins whose potency values are set to 100%. As multiple experimental values were available for many of the analogs, only the spread of mean values was considered to evaluate the accuracy of an FEP calculation. The spread of the mean values were captured by the width of the individual bar. The absolute spread including the experimentally reported errors are shown using the error-bar representations. A horizontal line in the bar represents the mean value (as mean of means).

In contrast to Insulin Wakayama and Chicago, the ΔΔG_bind_ obtained for insulin Los Angeles (-0.56 kcal/mol) was not in line with the experimental values (+1.54 kcal/mol) [Table S4]. Interestingly, in contrast to insulins Chicago and Wakayama, the PheB24Ser mutation that characterizes insulin Los Angeles retains a larger portion of its activity at the IR. The decomposition of ΔΔG_bind_ showed that change in electrostatics was nearly compensated by annihilation of phenyl ring atoms. Structural analysis revealed that the bulky B24Phe is buried in a hydrophobic pocket, which when mutated to a smaller hydrophilic serine residue could lead to partial or incomplete solvation of the sub-pocket that may cause the incorrect prediction of ΔΔG_bind_ (Figure 3c). To probe this aspect further, we reassessed the Serine end state by including one or two solvent molecules near the Serine residue. The position of solvent molecules near the residue was obtained by performing a short 50 ns MD simulations of the mutant. Structural analysis revealed that solvent molecules placed near the Serine residue moved away during the course of the calculation, indicating the complexities associated with solvation in this subpocket. Overall FEP could provide atomic insights into the loss of function associated with insulin Wakayama, Chicago, and more investigation is needed to understand insulin Los Angeles.

### Understanding the structural insights for the observed gain/loss-of-function associated with different classes of insulin analogs

To further understand receptor-ligand interactions, we aimed to quantify the effects of individual amino acid substitutions within the insulin molecule via evaluation of different classes of insulin analogs using FEP. Structural analysis accompanying the alchemical transformations performed with FEP was used to correlate structural drivers with the observed energy changes. For consistency, FEP calculations were performed using the same setup as used for evaluation of disease mutations. (**Figure 1, Figure S1a**).

Similar to the disease mutant calculations, no large fluctuations of structure were observed within the sphere during any of these simulations with RMSD values being < 0.3 Å. Once again, FEP schemes were devised for each analogue assessed in this study utilizing the fact that free energy is a state function and independent of path. In total 4.2 μs of simulations were performed, with an average of 420 ns used for each FEP calculation for each receptor. Here again, ΔΔG_bind_ with respect to WT-Ins or equivalent was used to quantify the relative affinities of each molecule to the IR in their respective conformation. In this work, insulins were divided into three classes, namely, commercially relevant analogs, and a series of mutations at the position B10, which has attracted a lot of attention due to the potential for increased potency, and truncated insulins. In total, 10 MD/FEP calculations covering 10 mutations were performed. Overall, FEP calculations could correctly capture the direction of shift in 9 out of 10 or in 90% of cases (Figure 4, Table S4). In seventy percent of these cases (7/10), the deviation between experiment and prediction was less than 1 kcal/mol (Figure 4, Table S4). As experimental functional profiles for analogs were in many cases obtained from multiple sources, the inner range (range of mean values) rather than the absolute spread of values including errors were considered to derive the mean of experimental ΔΔG_bind_, to assess the performance of FEP (Figure 4, Table S4).

### Commercially interesting Insulin analogs

Insulin analogs have been developed to specifically address different aspects of blood glucose control.^57,58^ In this study, we have performed FEP calculations of Insulin Lispro, Insulin Aspart and an insulin carrying a mutation A21gly, which is a metabolite (M1) of Insulin Glargine.^57,59,60^ Lispro is a fast-acting insulin analog from Eli Lilly.^57^ The sequence of Lispro differs from WT-Ins at the C-terminal end of the B-chain, involving a switch of residues B28Pro/B29Lys to B28Lys/B29Pro.^60,61^ Insulin Aspart carries a single amino-acid change from B28Pro to AspB28.^60^ Glargine is a long-acting insulin, carrying a glycine substitution at position A21 (Glargine metabolite M1), which we have characterized in this study, and the addition of two arginine molecules at the C-terminus of the B-chain (Glargine), whose effects have not been evaluated in this study, but is part of a related publication. The direction of shift in relative binding affinity (ΔΔG_bind_) values obtained from FEP calculations were in accordance with experiment in all three of the studied cases (100%) (Figure 4, Table S4).

For Aspart the calculated ΔΔG_bind_ value (-0.3 kcal/mol) recapitulated the near equipotency of Insulin aspart (∼0 kcal/mol) .^60,62–64^ Structural analysis revealed an interesting 90° rotation of the aromatic ring of B26Tyr residue in the presence of Asp at B28 (Figure 5) demonstrating the effect of this mutation on its neighbourhood. Similarly, the calculated value for lispro (+0.436 kcal/mol) was in good agreement with experiment (0.12 kcal/mol) [Table S4]. Decomposition of the energies revealed that the impact of electrostatics changes were not entirely compensated by the annihilation of Lysine side chains during the residue swap leading to the small magnitude of predicted free energy change. The energy analysis further showed that the proline transformations do not significantly impact the overall ΔΔG_bind_. Finally, the calculated ΔΔG_bind_ values for A21Gly (0.71 kcal/mol), showed a slightly exaggerated shift in potency when compared to near equipotency on Insulin Glargine M1 obtained from experiment (0.079 kcal/mol) [Table S4]. However, the calculated value was still within 1 kcal/mol of experiment and within experimental errors [Table S4]. Structural analysis supported a largely compensatory pattern of interactions when Asparagine or Glycine occupied the C-terminal residue of the insulin A-chain. On the whole, the Free Energy Perturbation (FEP) method precisely captured the energy profiles for all three commercially available insulin analogs, exhibiting a strong alignment with experimental results.

**Figure 5:**
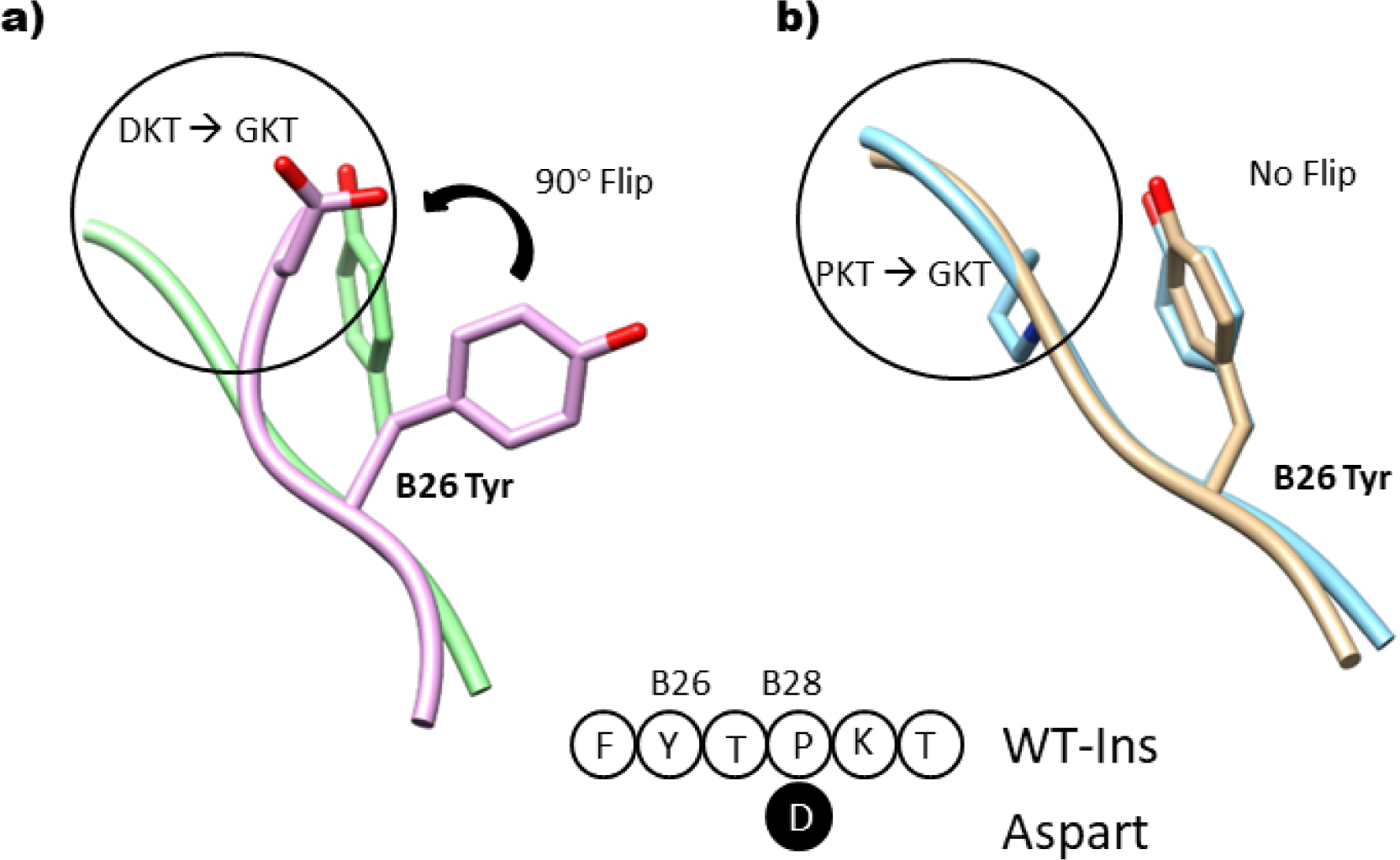
Analysis of Aspart showing the structural changes accompanying the transformation from WT-Ins to the mutant Aspart a) showing the flip of tyrosine residue in Aspart (magenta cartoon, B26Tyr and B28Asp shown as sticks), when compared to the glycine intermediate (green cartoon, B26Tyr and B28Gly shown as sticks) b) No flip of the tyrosine residue seen in WT-Ins (cyan cartoon, B26Tyr and B28Pro shown as sticks). This is again compared to the glycine intermediate (brown cartoon, B26Tyr and B28Gly shown as sticks). The sequence change accompanying the transformation of WT-Ins to Aspart is also shown.

### B10 Substitutions

In human insulin B10His plays a crucial role in the formation of insulin hexamers by coordinating with zinc.^65–67^ This position in the insulin sequence has also garnered significant interest as the substitution of the native histidine with an acidic residue (Asp or Glu) results in significant gain-of-function at the IR. ^58,68–70^ In the IR, B10His is seen proximal to an Arginine residue (510 from the FnIII-1’ domain) but the ND1 atom is still 4.5 Å away from the basic nitrogen (NH2) of the Arg. FEP was used to probe the effects of a variety of experimentally-evaluated substitutions at the B10 position to better understand correlations between the nature of amino acid at this position on overall function. Overall, FEP could accurately predict the direction of shift in all four cases (100%; Figure 4, Table S4).

FEP correctly captured the gain-of-function when Histidine at B10 is mutated to Asp (ΔΔG_bind_: -1.64 kcal/mol) albeit with an overestimation when compared to experiment (-0.45 kcal/mol) [Figure 4, Table S4]. Decomposition analysis revealed that a major portion of the gain in affinity towards the IR for B10Asp was driven by electrostatics which was only partially compensated by the annihilation of the imidazole ring atoms. Structural analysis of MD trajectories obtained from the simulation of B10Asp in complex with IR revealed that the Arg510 of FnIII-1’ forms a salt bridge when this position is occupied by an aspartic acid (Figure 6b). To further understand the molecular origins of the significant gain in potency at the IR we turned to solvent interaction network analysis from 50 ns MD simulations of the end states (B10His & B10Asp). This revealed a striking difference in the water mediated receptor ligand interactions between the WT-Ins and B10Asp at IR (Figure 6). The mutation of His to Asp at B10 resulted in large changes of first- and second-degree water mediated receptor-ligand contacts. First and second-degree solvent/solvent-mediated contacts are increased by ∼300% at the IR when B10 is occupied by aspartic acid compared to the native histidine (Figure 6, Table S5) further supporting the electrostatics-driven gain-of-function observed with this mutant.

**Figure 6:**
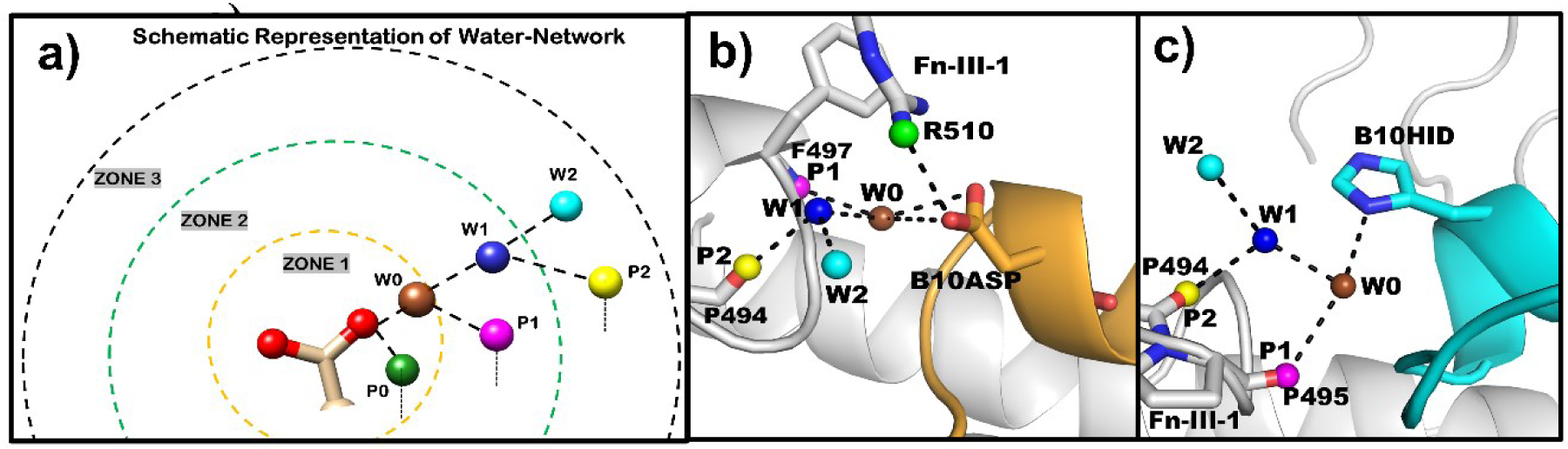
Analysis of the water-mediated interaction network around the position B10 of insulin in IR. A schematic representations of the classification of water-mediated interactions around a residue is shown. The Aspartic acid or Histidine residues occupying position B10 are shown in ball & stick representation, polar oxygen and nitrogen atoms are shown red or blue spheres respectively. First order or direct polar interactions with between B10 and a receptor residue is considered P0 and similarly W0 for a water contact. Second order contacts are classified as a W0-receptor polar contact defined as P1 or W0-water contact defined as W1. Similarly, third order contacts are defined as W1-receptor polar contact defined as P2 or W1-water contacts defined as W2. P0-2 and W0-2 are color coded as shown in the Figure (a) Shows the water mediated contacts for B10Asp-IR (b), B10His-IR (c). The figures highlight the increase in water-mediated polar interaction networks, when B10 is occupied by an Aspartic acid residue when compared to B10His in IR.

**Figure 7:**
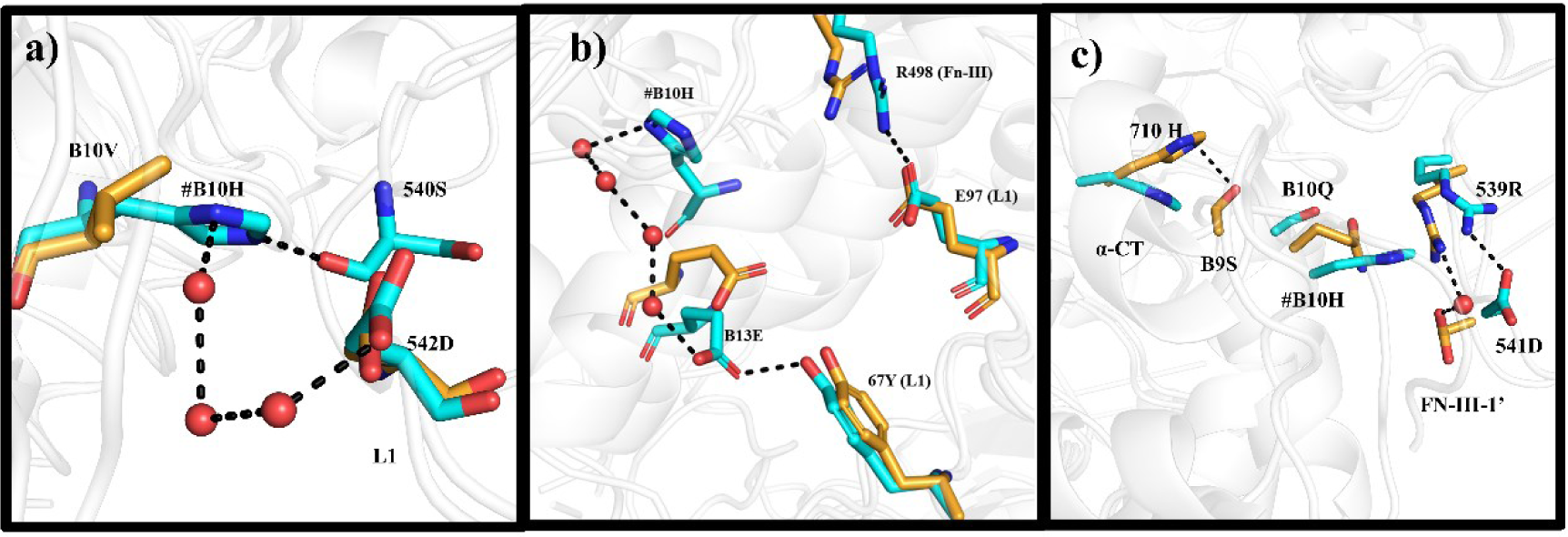
Structural changes of wildtype and a) B10Val, b) B10Ala, c) B10Gln were shown. The interacting residues in WT-Ins and mutant are shown as sticks with carbon atoms in cyan and orange respectively. Hydrogen bonds are shown as dashed lines. Water oxygen atoms are shown in red color spheres. WT-Ins was represented in cyan color; mutated state in orange color a) In WT-Ins, B10His has interactions with Serine from L1 domain and solvent mediated interactions, b) In B10Ala system, Interactions between 496Arg (FN-III-1’) and 97Glu were observed in WT-Ins, c) one-water mediated interaction between 539 and 541Asp were observed in B10Gln, additionally B9Ser interacts with 710His of α-CT domain.

In addition to B10Asp, Glendorf *et.al,* evaluated a series of ten different single amino acid substitutions at B10 position at IR, with respect to binding, metabolic and mitogenic potency.^65^ In this study, we have studied three additional representative amino acids mutations at B10, namely B10A (small), B10V, (hydrophobic alkyl), and B10Q (larger polar) to understand the effect of nature of amino acid on metabolic potencies. Whereas, their study considered DesB30-insulins, we performed the calculations with full-length insulin to maintain consistency, especially since a high-degree of correspondence between DesB30 and full-length insulins were reported.^71–75^ The calculated ΔΔG_bind_ values with full-length insulins for these analogs largely correlated with the observed experimental affinities at both the receptors (Figure 4, Table S4).

### B10Val & B10Ala

The substitution of histidine with an alanine or a small hydrophobic residue, such as valine, resulted in a sharp loss of function at the IR, which was accurately recapitulated by MD/FEP. In the case of B10Ala, the calculated ΔΔG_bind_ was +1.0 kcal/mol, whereas the experimental values was +0.18 kcal/mol [Table S4]. Structural analysis reveal that in WT-Ins, B10His interacts with Serine oxygen atom of L1-domain using the Nd atom of L1-domain, upon mutation with alanine this interaction is lost. In addition to the above interaction, in the mutated state the interactions between B13Glu and Tyr67 from the L1 domain is lost, pointing to this loss of interactions as the primary driver of the loss-of-function associated with this mutant. Similarly, FEP accurately captured the impact of substituting B10His with the larger hydrophobic residue Valine. The calculated ΔΔG_bind_ for B10Val is +0.29 kcal/mol whereas the experimental values for this mutation was +0.26 kcal/mol [Table S4]. Decomposition and structural analysis revealed a similar pattern to B10Ala where loss of interactions primarily drives the loss-of-function of the mutant when compared to WT-Ins.

### B10Gln

Mutation of B10 Histidine with a polar Glutamine (Q) is associated with a small gain of function. The ΔΔG_bind_ obtained from the FEP calculation accurately captured the gain in activity (-0.5 kcal/mol) when compared to the experimental value -0.018 kcal/mol.. Decomposition of ΔΔG_bind_ revealed that the change upon transformation of electrostatics was largely compensated by the annihilation of atoms, leading to a very subtly change in activity overall. Structural analysis revealed that B10Gln forms solvent mediated interaction that bridge 65Arg of L1-domain and 61Asp of Fn-III-1 domain. The 65Arg is also part of a salt-bridge with B13Glu of insulin, revealing the participation of B10Gln in an intricate polar interaction network around the B10 position. Overall, recapitulation of the effects of these different mutations at the crucial B10 position along with the analysis of the drivers of the observed changes from an energetic and structural viewpoint sheds light on the subtle interaction networks that govern Insulin-IR engagement in this neighbourhood.

### Truncated Insulins

Despite strong evolutionary conservation in the B-chain C-terminus for insulins, it was observed that the deletion of the residues (B26-B30) does not ablate its biological activity.^76–78^ Conversely, it was seen that the use of specific residues at the new C-terminus (B25) along with amidation created large increases in potency compared to WT-Ins.^76–78^ In this study, we evaluated three mutations at B25 for truncated [DesB26-30], WT-Ins, based on the work of Casaretto *et al*.^78^ The truncated mutants offered an opportunity to probe the generalizability of the FEP approach as deletion of the C-terminus represented a significant variation from WT-Ins. In-line with the experimental work, the C-terminus of the B-chain was amidated in our calculations.^77,78^ Overall, the FEP calculations could accurately capture direction of shift for two of three truncated systems (Figure 4, Table S4).

The calculated ΔΔG_bind_ for the mutation PheB25His (-1.5 kcal/mol) was in line with experiment (-0.7 kcal/mol). Structural analysis revealed the formation of a new network of water mediated, polar, and salt-bridge interactions between the receptor and ligand when histidine replaced phenylalanine as the C-terminus residue (Figure 8b). This enhanced interaction network was probably the driver of the increased potency observed with this mutation. Similarly, FEP correctly predicted the gain-of-function associated with the PheB25Tyr mutation (-1.99 kcal/mol FEP vs. -0.5 kcal/mol from experiment). Decomposition of ΔΔG_bind_ revealed the significant change in electrostatics was not compensated by annihilation of the hydroxyl group of the Tyrosine (Mutant). Structural analysis supported the gain in electrostatics upon mutating Phe to Tyr and revealed a new interaction of B25Tyr with the Glutamine (N15) of the L1-domain. Unlike, the previous two cases FEP failed to predict the loss-of-function associated with PheB25Trp (Figure 4, Table S4). Decomposition of interactions have shown that a majority of the ΔΔG_bind_ originates upon the annihilation of the distal aromatic carbon atoms in the tryptophan, which may be due to incomplete reorganization of the interaction network following such a large perturbation. Overall, Free Energy Perturbation (FEP) analysis successfully captured two out of three mutations in truncated insulin systems. Analyzing the underlying factors behind these observed changes from both an energetic & structural perspective sheds light on the dynamics of truncated insulin-IR interactions and offered clues as to how improved activity is possible, despite large-scale deletions.

**Figure 8:**
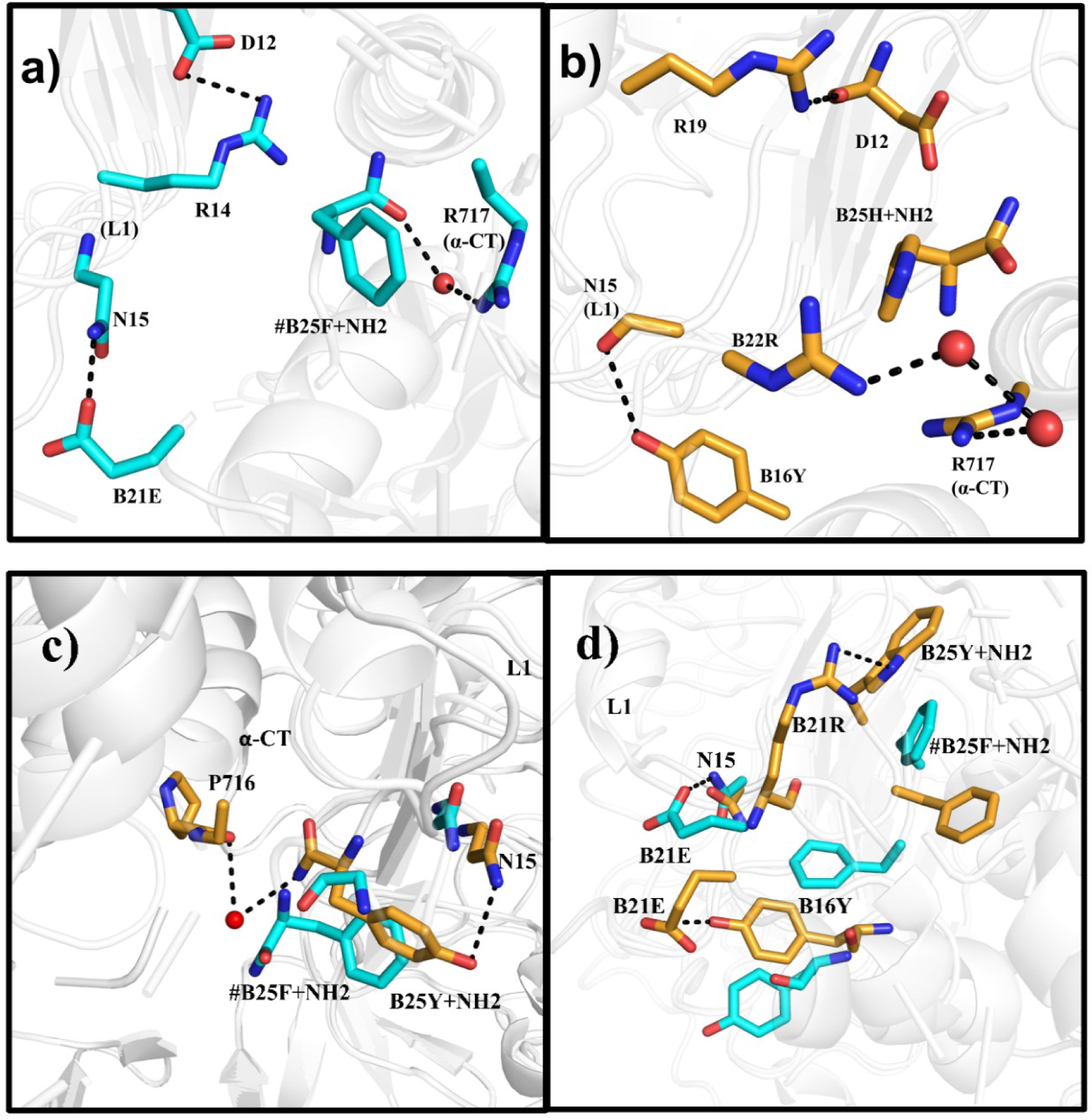
Structural analysis of truncated insulins (desB26-B30) for the three systems considered were shown. In a) Truncated insulin with C-terminus occupied by the natural WT residue i.e. B25Phe is shown. In general, the receptor-ligand complex is shown as a cartoon with carbon atoms in cyan. The highlighted interaction networks are shown with participating residues in sticks with carbon atoms in cyan. b) Truncated insulin with C-terminus occupied by the mutant B25His is shown in orange stick representation. c) Truncated B25Tyr insulin is shown, where B15Tyr forms interactions with 15Asn of L1 domain. d) Truncated B25Trp insulin is shown, where B21Arg forms interactions with Tryptophan and B16Tyr forms interaction with Glutamic acid. In general, the receptor-ligand complex is shown as a cartoon with carbon atoms in grey. The highlighted interaction networks are shown with participating residues in sticks with carbon atoms in orange.

## Discussion

A century on from the discovery of insulin, molecular-level understanding of its receptor engagement remains an important goal.^2,15,21,79^ In this work, we have leveraged the recent increase in high-resolution structures of the IR^18,25,28,29,51^ and used MD/FEP to elucidate atomistic drivers of receptor-ligand engagement. Overall, thirteen systems were examined utilizing MD/FEP and good agreement with experiment was observed in ∼85% of the cases. Consequently, three key results emerged from our evaluations. First, MD simulations captured the stability of bound insulin in the HA site of the IR and revealed key long-lived interactions of insulin with IR. This provided us with a framework to evaluate and understand insulin recognition at its cognate receptor. Next, FEP could capture the loss of function for two disease-causing mutations, namely Insulin Wakayama and Insulin Chicago. Decomposition of the free energy changes associated with the disease-causing mutations and structural analysis of the simulations provided atomistic explanations for the loss-of-function at the IR. Finally, FEP managed to accurately recapitulate the efficacy profiles for a variety of man-made insulin variants including molecules with significant changes in sequence length. Additionally, these calculations provided a much greater depth of understanding of the link between molecular-level drivers and the observed functional profiles of these different insulin variants.

Recent structural breakthroughs have resulted in high-resolution structures of IR and this served as a starting point for MD simulations of the insulin-IR complex. ^2,18,25–29,51,80,81^ The short MD simulations confirmed the global stability of insulin-IR complex with regards to the HA site. Analysis of close, long-lived contacts, defined as a 4.5 Å sphere around the ligand, re-emphasized the importance of the α-CT in ligand recognition, which is in-line with previous studies. Further, the observed long-lived interactions, also captured previously known key residues involved in receptor binding such as A19Tyr .^25,82,83^ Our analysis also revealed the importance of glycine residues, such as B8 and B23 in terms of maintaining the bound conformation of insulin.^28,83^ It was further observed that the valine at B12, which is part of the structured helical domain of the B-chain was involved in stable interactions with 714Phe of the α-CT, which was in-line with previous results, showing significant loss-of-function upon mutation of this residue to almost any other amino acid.^70^ In a recent mutational investigation conducted by Kertisova *et al.,* the pivotal significance of 717Arg within the α-CT domain was underscored. This residue, in conjunction with other residues participating in the transition from the apo-form to the bound-form, plays a crucial role. Any substitution of this residue with different amino acids results in inactivation, and this interaction was captured through molecular dynamics (MD) simulations.^84^ Overall, these simulations provided us with a global understanding of the patterns of recognition of human insulin at the IR.

Whereas, the short MD simulations provided us with a broad picture of insulin recognition at IR, understanding the changes in functional profile caused by mutation of individual residues is key to developing an enhanced understanding of receptor-ligand engagement. As a first step towards this we wished to study the three known naturally occurring point mutations of human insulin that increase the likelihood of developing diabetes.^53,55^ The three variants, named Wakayama (ValA3Leu), Los Angeles (PheB24Ser), and Chicago (PheB25Leu) result in loss-of-function at the IR.^53,55^ An illustration of the power of MD/FEP was seen in the evaluation of Insulin Wakayama. Whereas, Valine to Leucine appears as a conservative mutation, there is a significant loss-of-function associated with this change to WT-Ins. From the calculations, a disruption of direct and solvent-mediated interactions between the insulin variant and the L1 domain of the IR was revealed, which was also reflected in the unfavorable free energy change associated with this mutation. Similarly, a loss-of-function was predicted by our calculation for the Chicago variant, where a similar disruption of interaction networks between the receptor and ligand were identified. Interestingly, B24Ser that characterizes the Los Angeles variant has been shown to be one of the mutations that still retains a reasonable degree of metabolic potency .^85,86^ Our calculations indicated a small increase in potency at the IR, and did not capture the loss-of-function associated with this mutant. B24Phe resides in a buried hydrophobic pocket and mutation to the smaller hydrophilic Serine will probably result in a partial solvation of this pocket. The time-scales for our FEP calculations may not have been sufficient to allow for the required solvation of this buried pocket. Hence, to better understand the above result, we performed MD simulations in presence of a Serine at B24 and allowed for partial solvation of this pocket. However, the use of a partially solvated topology in the presence of serine also did not drastically alter the results. Interestingly, structural analysis of the simulations revealed that the solvent molecules in the pocket move away from Serine at B24 during the calculation. Hence, more work is required to better understand the interaction patterns, when a small polar amino acid such as Serine occupies the B24Phe sub-pocket. Additionally, since our work was restricted to the HA binding site of insulin, we would also be looking at the LA site in future work to evaluate if disruption of binding there was driving the functional profile of this disease variant.

In order to further probe the contributions of individual amino acids within insulin towards potency via the IR, we quantified the free energy changes associated with man-made mutations using FEP. In the last several decades, a number of insulin variants have been produced and characterized.^57,60,62,87–89^ Two such approved insulin analogs Lispro and Aspart harbor changes at the C-terminal end of the insulin B-chain.^57,60,62^ The FEP calculations accurately captured the near equipotency of Lispro and Aspart to WT-Ins. Structural analysis also revealed how mutations at one position in the sequence affects the conformations of neighbouring residues as seen in the case of Aspart. Structural and solvent-network analysis around these residues further confirmed significant reorganization of water networks occurred around the sites of annihilation and showed how these perturbations can often be compensatory. Another commercially interesting variant, the A21Gly mutant (a metabolite of Insulin Glargine: M1) was predicted by MD-FEP to be nearly equipotent at IR compared to WT-Ins with minimal structural perturbation.^59^ Overall, the accuracy of the methods in capturing the functional profiles of the three commercially interesting insulin analogs considered in this study showed great promise for its use in insulin design and allowed for a better understanding of the role of the two C-termini with respect to IR engagement.

The B10 position within the insulin sequence has attracted significant attention due to the super-potency (∼375% of human insulin) associated with the mutation of the natural histidine at this position to acidic residues such as aspartic or glutamic acid.^62,65,66,66,71,90^ Hence, understanding the structural drivers of the potency increase could prove important in leveraging the potential of B10 mutants.^65^ The MD-FEP calculations managed to capture the significant increase in potency associated with the B10Asp variant at the IR. Structural analysis at the IR, revealed the importance of interaction with R510 of FnIII-1ˈ domain on the increased potency of B10Asp. Additionally, analysis of the solvent networks around the residue, further revealed that the change from histidine to aspartic acid promoted a strong network of water-mediated interactions including inter-domain solvent bridges that stabilized the overall bound conformation at the IR. The solvent network analysis demonstrates the power of MD simulations, where the use of explicit water molecules can capture important effects not readily apparent from static structures or even continuum solvation methods. The FEP could also capture the loss of function for hydrophobic residues Valine, Alanine, where the change in microenvironment when B10His is replaced with non-polar amino acids changes lead to loss of interaction between insulin and L1-domain. The subtle increase in potency when a glutamine occupies B10Gln was also captured by our calculation. A network of polar and solvent mediated interactions mediated by B10Gln was shown to be a driver of the marginal gain-in-function for this variant. The insights into the molecular drivers of the observed changes when this position is altered could be key in taking advantage of B10 variants to create novel insulins with tailored functional profiles.

Previous studies have shown that deleting the last five highly conserved residues at the C-terminus of the insulin B-chain still results in a molecule with significant potency.^77,78,91^ In fact, changes in the new C-terminus (B25) of these truncated molecules, provide variants with increased potency at the IR compared to WT-Ins.^77,78,92^ These insulins of different length and C-terminal modifications provided an ideal opportunity to explore the generalizability of MD/FEP as an *in silico* mutagenesis tool in peptide design. Despite, truncation and amidation of the B-chain C-terminus, FEP was able to accurately reproduce the experimental functional profiles in two of three cases. The increased potency associated with Histidine and Tyrosine as C-termini was captured by our calculations along with an understanding of its molecular origins, namely an altered polar interaction network around the new residues. The FEP could not capture the loss-of-function associated with Tryptophan as C-termini. This may be attributed to insufficient time-scales required to capture the changes in microenviroment during the annihilation of the large tryptophan group.

## Conclusion

In this study, we have demonstrated the utility of MD/FEP in accurately capturing and elucidating the molecular drivers for the loss/gain-of-function for a variety of insulin analogs at IR. This consistency was noticed across different positions within the insulin sequence, spanning a wide range of amino acid mutations, and also encompassed variants with significant changes in sequence length and amidation of the C-terminus. The recapitulation of the functional profiles was accompanied by an ability to deconvoluate the free energy changes into individual energetic contributions and rationalize these from a structural perspective. In summary, this study has created an enhanced understanding of insulin recognition at the IR and showcased the power of MD/FEP as a generalized *in silico* mutation evaluation tool for the rational design of novel insulins and peptides in general.

## Methods

### Homology modelling

Homology modelling (comparative modelling) is used to predict three-dimensional structure of protein from the amino acid sequences.^93^ In this method, a closely resembled protein or protein from similar family is used as template to determine the structure of the other protein (query) for which sequence is known. This method frequently used to determine the structure of missing domains, stretch of residues in loop regions of the protein obtained from protein data band (PDB) from various experiments. In this study, we have used Modeller 9.25 version software for homology modelling missing residues in IR (generally few residues in loop regions).^52,94^ The final structure was selected based on the DOPE (discrete optimized protein energy) score and visual inspection.

### Molecular Dynamic (MD) Simulations

The MD simulations were performed with the initial coordinates obtained from experimental structures for IR cognate complexes with PDB accession code: 6CE9.^45,51^ The missing residues/segments were modelled using available templates in Modeller software version 9.25.^52^ The missing residues were mostly at the terminal loop regions of the domains in the structures considered. Protonation states of the amino acids in the protein has effect on the overall binding energy of the ligand.^95^ Therefore careful choice of protonation states are crucial to obtain relevant binding energy values. The protonation state for ionizable residues were set to their most probable state ^96,97^ and histidine tautomers were assigned by visual inspection. The selected models were fully solvated using TIP3P^98^ water model and counter ions were added to neutralize the system. The neutral solvated system is minimized and then performed equilibration in NPT ensemble for 10 ns using AMBER14ffSB^99^ force field in AMBER^100^. During equilibration, restraints were placed on heavy atoms of protein allowing solvent to adjust around the protein, which then were removed in a series of equilibration steps spanning a total of 10 ns. After equilibrating the system, a production run of 50 ns was performed at 300 K temperature with a non-bonded cut-off of 10 Å. The SHAKE algorithm was used to constrain bond lengths during the simulation.^101^ Long range electrostatics were treated with Ewald summation method.^102^ The Langevin thermostat was used to maintain the temperature on the system, while Berendsen barostat maintained pressure at 1 Bar with a coupling of 2 ps.^103,104^ The final structure obtained after 10 ns equilibration from the MD simulations using periodic boundary conditions is used as input to compute free energy (ΔΔG) using FEP method with spherical boundary conditions (SBC).

### MD using SBC and the Free Energy Perturbation (FEP) Method

The free energy (ΔΔG_bind_) were computed using FEP implemented in Q program.^105^ The final snapshot from the equilibration was used as starting structure for the FEP calculations using spherical boundary conditions. In spherical boundary conditions (SBC), the region of interest or the active site is solvated, whereas the rest of the biomolecular system remains outside. A representative image for conversion of periodic boundary conditions (PBC) to spherical boundary conditions is provided in the supplementary information (Figure S1a). The calculations using SBC conditions are faster compared to PBC as the number of solvent molecules, receptor atoms are lesser and treatment of charges are well taken in comparison to PBC. In the current study, we have chosen Cδ atom of B15Leu of insulin as center and considered a sphere with radius 27 Å at IR. Heavy restraints were applied for the atoms outside the sphere in order to maintain the structure close to the initial structure.

All the protein, ligand and water atoms with the defined radius are explicitly considered. OPLSAA force field were used for the receptor complex. Thus obtained system was equilibrated in a series of six steps with varying temperature and restraints on the heavy atoms. Finally, the well equilibrated system was used as input to run unrestrained production run which was used to compute free energies. Atoms close to sphere edge were restrained to their initial coordinates and atoms beyond sphere edge were excluded from nonbonded interactions. Asp, Glu, Lys and Arg residues within radius of 27 Å are protonated according to their most probable state at pH 7 and ionizable residues close to sphere were neutralized. Residues lying outside the sphere are restrained to stay close to the observed structure, so that we can evaluate energy of this conformational region rather than allowing system to move into other conformational regions.^106^ The protonation state for histidine’s within sphere were assigned based on visual inspection. The SHAKE algorithm was used to constraint all the bonds, all angles and water molecules at the sphere surface were subjected to radial and polarization restraints according to the SCAAS model.^101,104,107^ Long range electrostatics were treated with local reaction field (LRF) method.^108^ The time step was set to 1 fs and non-bonded pair lists were updated every 25 steps.

The following FEP protocol was carried out to obtain relative binding free energy for the mutants: i) the transformation of partial charges and ii) combined transformation of Lennard-Jones (LJ) and parameters involving covalent bonds in several MD/FEP calculations. To annihilate heavy atoms, a separate MD/FEP calculations were performed to remove them in step-wise manner. A softcore potential was introduced for the atom in a first step and followed by removal of the resulting van der Waal’s potential. The force field parameters describing angles, bonds, and proper/improper dihedrals were retained for annihilated atoms. Each FEP calculation were divided into *n* intermediate states which were equilibrated parallely. The potential (U_m_) for the respective transformation A (initial state) and B (final state) is given by

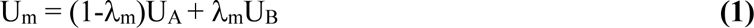

where λ_m_ varies from 0 to 1. A value of λ_m_=0 represents initial state, λ_m_=1 represents final state and values in between 0 and 1 represents intermediate states. The transformations involving partial charges were performed using up to 36 λ_m_ steps and the number of λ_m_ steps used to transform LJ and bonded parameters ranged from 42 to 63 λ_m_ steps depending upon the complexity of the transformation. The ligand-receptor complex was typically equilibrated for 1.76 ns at each λ_m_ step. During the equilibration of the system the harmonic restraints were released on ligand-receptor complexes in several steps and the temperature was increased in a series of steps to 310 K. After equilibration, the unrestrained simulation for the complex was typically carried out for 0.5 ns for each λ_m_ step from which potential energies were extracted.

The ligand in water consisted of the entire insulin molecule and was typically equilibrated for 0.76 ns followed by 0.25 ns of unrestrained simulation for each λ_m_ step. The same sphere radii were maintained between the respective receptor and water simulations. The difference in free energy between initial state (A) and final state (B) was calculated by summing up the free energy differences of the n intermediate states using the relation

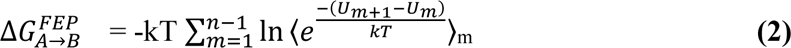

Where 〈… 〉_*m*_ represents an ensemble average on the potential U_m_ which is calculated from the MD simulations, k is Boltzmann constant and T is absolute temperature.^109^ The uncertainty of a transformation was quantified as the difference in free energy obtained by applying FEP formula in the forward and reverse direction and was minimized by increasing the number of λm values or simulation length until convergence was obtained.

### Water(solvent)-interaction network analysis

To assess the solvent and solvent-mediated interaction networks around a selected residue a distance-based analysis scheme was devised. Initially, snapshots from the MD trajectories were saved as PDB structures. Hydrogens atoms were removed from the snapshots obtained from the MD trajectories and the residue of interest was provided as an input. Only side chain heteroatoms of the defined residue on interest were considered for the analysis. Polar contacts with either water or protein were defined within a 3.5 Å radius. First order contacts are those polar contacts with the receptor that were within 3.5 Å of the residue of interest (P0), and waters within the same cutoff (W0). Using the W0 waters as centers, the analysis was repeated. A receptor polar contact within 3.5 Å of any W0 water was defined as a second order contact (P1) and any water molecule within the same radius as a second-order water contact (W1), The process was repeated with W1 waters as centers resulting in a list of third-order receptor (P2) or water (W2) contacts. Average values of W0, P0, W1, P1, W2 and P2 for any residue of interest were obtained by averaging across the number of considered snapshots.

## Acknowledgments

We thank Sudha Ravishankar, Samdani Ansar and Sampath Ranganathan for their valuable inputs in the preparation of this manuscript.

## Author Contributions

M.M.S., R.C., and A.R. designed the research project. M.M.S., R.C., and A.R. performed and analysed the MD simulations and FEP calculations. All authors contributed to the subsequent discussion of the results. All authors contributed to the writing of the manuscript.

## References

(1) Saltiel, A. R.; Kahn, C. R. Insulin Signalling and the Regulation of Glucose and Lipid Metabolism. Nature 2001, 414 (6865), 799–806.

(2) Gutmann, T.; Schäfer, I. B.; Poojari, C.; Brankatschk, B.; Vattulainen, I.; Strauss, M.; Coskun, Ü. Cryo-EM Structure of the Complete and Ligand-Saturated Insulin Receptor Ectodomain. Journal of Cell Biology 2019, 219 (1), e201907210.

(3) Rahman, M. S.; Hossain, K. S.; Das, S.; Kundu, S.; Adegoke, E. O.; Rahman, Md. A.; Hannan, Md. A.; Uddin, M. J.; Pang, M.-G. Role of Insulin in Health and Disease: An Update. Int J Mol Sci 2021, 22 (12), 6403.

(4) Wilcox, G. Insulin and Insulin Resistance. Clin Biochem Rev 2005, 26 (2), 19–39.

(5) Tatulian, S. A. Structural Dynamics of Insulin Receptor and Transmembrane Signaling. Biochemistry 2015, 54 (36), 5523–5532.

(6) Bentley, G.; Dodson, E.; Dodson, G.; Hodgkin, D.; Mercola, D. Structure of Insulin in 4-Zinc Insulin. Nature 1976, 261 (5556), 166–168.

(7) Smith, G. D.; Swenson, D. C.; Dodson, E. J.; Dodson, G. G.; Reynolds, C. D. Structural Stability in the 4-Zinc Human Insulin Hexamer. Proceedings of the National Academy of Sciences 1984, 81 (22), 7093–7097.

(8) Ciszak, E.; Beals, J. M.; Frank, B. H.; Baker, J. C.; Carter, N. D.; Smith, G. D. Role of C-Terminal B-Chain Residues in Insulin Assembly: The Structure of Hexameric LysB28ProB29-Human Insulin. Structure 1995, 3 (6), 615–622.

(9) Busto-Moner, L.; Feng, C.-J.; Antoszewski, A.; Tokmakoff, A.; Dinner, A. R. Structural Ensemble of the Insulin Monomer. Biochemistry 2021, 60 (42), 3125–3136.

(10) Steiner, D. F.; Oyer, P. E. The Biosynthesis of Insulin and a Probable Precursor of Insulin by a Human Islet Cell Adenoma. Proceedings of the National Academy of Sciences 1967, 57 (2), 473–480.

(11) Steiner, Donald F., Dennis Cunningham, Lilian Spigelman, and Bradley Aten. “Insulin biosynthesis: evidence for a precursor.” Science 157, no. 3789 (1967): 697–700. *Insulin*

(12) Vasiljević, J.; Torkko, J. M.; Knoch, K.-P.; Solimena, M. The Making of Insulin in Health and Disease. Diabetologia 2020, 63 (10), 1981–1989.

(13) Arvan, P.; Halban, P. A. Sorting Ourselves Out: Seeking Consensus on Trafficking in the Beta-Cell. Traffic 2004, 5 (1), 53–61.

(14) Dodson, G.; Steiner, D. The Role of Assembly in Insulin’s Biosynthesis. Current Opinion in Structural Biology 1998, 8 (2), 189–194.

(15) De Meyts, Pierre. “Insulin and its receptor: structure, function and evolution.” Bioessays 26, no. 12 (2004): 1351–1362.

(16) Mayer, J. P.; Zhang, F.; DiMarchi, R. D. Insulin Structure and Function. Peptide Science 2007, 88 (5), 687–713. 10.1002/bip.20734.

(17) Chang, Seung-Gu, Ki-Doo Choi, Seung-Hwan Jang, and Hang-Cheol Shin. “Role of disulfide bonds in the structure and activity of human insulin.” Molecules & Cells (Springer Science & Business Media BV) 16, no. 3 (2003).

(18) Menting, John G., Jonathan Whittaker, Mai B. Margetts, Linda J. Whittaker, Geoffrey K-W. Kong, Brian J. Smith, Christopher J. Watson et al. “How insulin engages its primary binding site on the insulin receptor.” Nature 493, no. 7431 (2013): 241–245.

(19) Papaioannou, A.; Kuyucak, S.; Kuncic, Z. Computational Study of the Activity, Dynamics, Energetics and Conformations of Insulin Analogues Using Molecular Dynamics Simulations: Application to Hyperinsulinemia and the Critical Residue B26. Biochemistry and Biophysics Reports 2017, 11, 182–190.

(20) Žáková, L.; Kletvíková, E.; Veverka, V.; Lepšík, M.; Watson, C. J.; Turkenburg, J. P.; Jiráček, J.; Brzozowski, A. M. Structural Integrity of the B24 Site in Human Insulin Is Important for Hormone Functionality *. Journal of Biological Chemistry 2013, 288 (15), 10230–10240.

(21) Mathieu, Chantal, Pieter-Jan Martens, and Roman Vangoitsenhoven. “One hundred years of insulin therapy.” Nature Reviews Endocrinology 17, no. 12 (2021): 715–725.

(22) De Meyts, P.; Whittaker, J. Structural Biology of Insulin and IGF1 Receptors: Implications for Drug Design. Nat Rev Drug Discov 2002, 1 (10), 769–783.

(23) McKern, N. M.; Lawrence, M. C.; Streltsov, V. A.; Lou, M.-Z.; Adams, T. E.; Lovrecz, G. O.; Elleman, T. C.; Richards, K. M.; Bentley, J. D.; Pilling, P. A.; Hoyne, P. A.; Cartledge, K. A.; Pham, T. M.; Lewis, J. L.; Sankovich, S. E.; Stoichevska, V.; Da Silva, E.; Robinson, C. P.; Frenkel, M. J.; Sparrow, L. G.; Fernley, R. T.; Epa, V. C.; Ward, C. W. Structure of the Insulin Receptor Ectodomain Reveals a Folded-over Conformation. Nature 2006, 443 (7108), 218–221.

(24) Virkamäki, A.; Ueki, K.; Kahn, C. R. Protein–Protein Interaction in Insulin Signaling and the Molecular Mechanisms of Insulin Resistance. J Clin Invest 1999, 103 (7), 931–943.

(25) Scapin, Giovanna, Venkata P. Dandey, Zhening Zhang, Winifred Prosise, Alan Hruza, Theresa Kelly, Todd Mayhood, Corey Strickland, Clinton S. Potter, and Bridget Carragher. “Structure of the insulin receptor–insulin complex by single-particle cryo-EM analysis.” Nature 556, no. 7699 (2018): 122–125.

(26) Uchikawa, E.; Choi, E.; Shang, G.; Yu, H.; Bai, X. Activation Mechanism of the Insulin Receptor Revealed by Cryo-EM Structure of the Fully Liganded Receptor–Ligand Complex. eLife 2019, 8, e48630.

(27) Nielsen, J.; Brandt, J.; Boesen, T.; Hummelshøj, T.; Slaaby, R.; Schluckebier, G.; Nissen, P. Structural Investigations of Full-Length Insulin Receptor Dynamics and Signalling. Journal of Molecular Biology 2022, 434 (5), 167458.

(28) Menting, J. G.; Yang, Y.; Chan, S. J.; Phillips, N. B.; Smith, B. J.; Whittaker, J.; Wickramasinghe, N. P.; Whittaker, L. J.; Pandyarajan, V.; Wan, Z.; Yadav, S. P.; Carroll, J. M.; Strokes, N.; Roberts, C. T.; Ismail-Beigi, F.; Milewski, W.; Steiner, D. F.; Chauhan, V. S.; Ward, C. W.; Weiss, M. A.; Lawrence, M. C. Protective Hinge in Insulin Opens to Enable Its Receptor Engagement. Proc Natl Acad Sci U S A 2014, 111 (33), E3395–E3404.

(29) Croll, T. I.; Smith, B. J.; Margetts, M. B.; Whittaker, J.; Weiss, M. A.; Ward, C. W.; Lawrence, M. C. Higher-Resolution Structure of the Human Insulin Receptor Ectodomain: Multi-Modal Inclusion of the Insert Domain. Structure 2016, 24 (3), 469–476.

(30) Baker, Edward N., Thomas Leon Blundell, John F. Cutfield, Eleanor Joy Dodson, George Guy Dodson, Dorothy Mary Crowfoot Hodgkin, Roderick E. Hubbard et al. “The structure of 2Zn pig insulin crystals at 1.5 Å resolution.” Philosophical Transactions of the Royal Society of London. B, Biological Sciences 319, no. 1195 (1988): 369–456.

(31) Olsen, H. B.; Ludvigsen, S.; Kaarsholm, N. C. Solution Structure of an Engineered Insulin Monomer at Neutral PH. Biochemistry 1996, 35 (27), 8836–8845.

(32) A. Hua, Q.-X.; Gozani, S. N.; Chance, R. E.; Hoffmann, J. A.; Frank, B. H.; Weiss, M. A. Structure of a Protein in a Kinetic Trap. Nat Struct Mol Biol 1995, 2 (2), 129–138..

(33) Gorai, B.; Vashisth, H. Progress in Simulation Studies of Insulin Structure and Function. Front Endocrinol (Lausanne) 2022, 13, 908724.

(34) Papaioannou, A.; Kuyucak, S.; Kuncic, Z. Molecular Dynamics Simulations of Insulin: Elucidating the Conformational Changes That Enable Its Binding. PLOS ONE 2015, 10 (12), e0144058.

(35) Kim, Y. H.; Kastner, K.; Abdul-Wahid, B.; Izaguirre, J. A. Evaluation of Conformational Changes in Diabetes-Associated Mutation in Insulin a Chain: A Molecular Dynamics Study. *Proteins: Structure*, Function, and Bioinformatics 2015, 83 (4), 662–669.

(36) Bagchi, K.; Roy, S. Sensitivity of Water Dynamics to Biologically Significant Surfaces of Monomeric Insulin: Role of Topology and Electrostatic Interactions. J. Phys. Chem. B 2014, 118 (14), 3805–3813.

(37) Mark, A. E.; Berendsen, H. J. C.; Gunsteren, W. F. V. Conformational flexibility of aqueous monomeric and dimeric insulin: a molecular dynamics study. ACS Publications.

(38) Schlitter, J.; Engels, M.; Krüger, P.; Jacoby, E.; Wollmer, A. Targeted Molecular Dynamics Simulation of Conformational Change-Application to the T ↔ R Transition in Insulin. Molecular Simulation 1993, 10 (2–6), 291–308.

(39) Zoete, V.; Meuwly, M.; Karplus, M. Study of the Insulin Dimerization: Binding Free Energy Calculations and per-Residue Free Energy Decomposition. Proteins: Structure, Function, and Bioinformatics 2005, 61 (1), 79–93.

(40) Zoete, V.; Meuwly, M. Importance of individual side chains for the stability of a protein fold: Computational alanine scanning of the insulin monomer. Journal of Computational Chemistry 2006, 27 (15), 1843–1857.

(41) Todorova, N.; Marinelli, F.; Piana, S.; Yarovsky, I. Exploring the Folding Free Energy Landscape of Insulin Using Bias Exchange Metadynamics. J. Phys. Chem. B 2009, 113 (11), 3556–3564.

(42) Kim, T.; Rhee, A.; Yip, C. M. Force-Induced Insulin Dimer Dissociation: A Molecular Dynamics Study. J. Am. Chem. Soc. 2006, 128 (16), 5330–5331.

(43) Kiselyov, Vladislav V., Soetkin Versteyhe, Lisbeth Gauguin, and Pierre De Meyts. “Harmonic oscillator model of the insulin and IGF1 receptors’ allosteric binding and activation.” Molecular systems biology 5, no. 1 (2009): 243.

(44) Vashisth, H.; Abrams, C. F. Docking of insulin to a structurally equilibrated insulin receptor ectodomain. Proteins: Structure, Function, and Bioinformatics 2010, 78 (6), 1531–1543.

(45) Scapin, G.; Dandey, V. P.; Zhang, Z.; Prosise, W.; Hruza, A.; Kelly, T.; Mayhood, T.; Strickland, C.; Potter, C. S.; Carragher, B. Structure of the Insulin Receptor-Insulin Complex by Single-Particle Cryo-EM Analysis. Nature 2018, 556 (7699), 122–125..

(46) Vanga, S. R.; Åqvist, J.; Hallberg, A.; Gutiérrez-de-Terán, H. Structural Basis of Inhibition of Human Insulin-Regulated Aminopeptidase (IRAP) by Benzopyran-Based Inhibitors. Frontiers in Molecular Biosciences 2021, 8.

(47) Wang, L.; Wu, Y.; Deng, Y.; Kim, B.; Pierce, L.; Krilov, G.; Lupyan, D.; Robinson, S.; Dahlgren, M. K.; Greenwood, J.; Romero, D. L.; Masse, C.; Knight, J. L.; Steinbrecher, T.; Beuming, T.; Damm, W.; Harder, E.; Sherman, W.; Brewer, M.; Wester, R.; Murcko, M.; Frye, L.; Farid, R.; Lin, T.; Mobley, D. L.; Jorgensen, W. L.; Berne, B. J.; Friesner, R. A.; Abel, R. Accurate and Reliable Prediction of Relative Ligand Binding Potency in Prospective Drug Discovery by Way of a Modern Free-Energy Calculation Protocol and Force Field. J. Am. Chem. Soc. 2015, 137 (7), 2695–2703.

(48) Turvey, S. J.; McPhillie, M. J.; Kearney, M. T.; Muench, S. P.; Simmons, K. J.; Fishwick, C. W. G. Recent Developments in the Structural Characterisation of the IR and IGF1R: Implications for the Design of IR–IGF1R Hybrid Receptor Modulators. RSC Med Chem 13 (4), 360–374.

(49) Sindhikara, D.; Wagner, M.; Gkeka, P.; Güssregen, S.; Tiwari, G.; Hessler, G.; Yapici, E.; Li, Z.; Evers, A. Automated Design of Macrocycles for Therapeutic Applications: From Small Molecules to Peptides and Proteins. J. Med. Chem. 2020, 63 (20), 12100–12115.

(50) Zhang, X.; Yu, D.; Sun, J.; Wu, Y.; Gong, J.; Li, X.; Liu, L.; Liu, S.; Liu, J.; Wu, Y.; Li, D.; Ma, Y.; Han, X.; Zhu, Y.; Wu, Z.; Wang, Y.; Ouyang, Q.; Wang, T. Visualization of Ligand-Bound Ectodomain Assembly in the Full-Length Human IGF-1 Receptor by Cryo-EM Single-Particle Analysis. Structure 2020, 28 (5), 555–561.e4.

(51) Xu, Y.; Kong, G. K.-W.; Menting, J. G.; Margetts, M. B.; Delaine, C. A.; Jenkin, L. M.; Kiselyov, V. V.; De Meyts, P.; Forbes, B. E.; Lawrence, M. C. How Ligand Binds to the Type 1 Insulin-like Growth Factor Receptor. Nat Commun 2018, 9 (1), 821.

(52) Eswar, N.; Webb, B.; Marti-Renom, M. A.; Madhusudhan, M. s.; Eramian, D.; Shen, M.; Pieper, U.; Sali, A. Comparative Protein Structure Modeling Using Modeller. Current Protocols in Bioinformatics 2006, 15 (1), 5.6.1–5.6.30.

(53) Nishi, M.; Nanjo, K. Insulin Gene Mutations and Diabetes. Journal of Diabetes Investigation 2011, 2 (2), 92–100.

(54) Tager, H., B. Given, D. Baldwin, M. Mako, J. Markese, A. Rubenstein, J. Olefsky, M. Kobayashi, O. Kolterman, and R. Poucher. “A structurally abnormal insulin causing human diabetes.” Nature 281, no. 5727 (1979): 122–125.

(55) Shoelson, S.; Haneda, M.; Blix, P.; Nanjo, A.; Sanke, T.; Inouye, K.; Steiner, D.; Rubenstein, A.; Tager, H. Three Mutant Insulins in Man. Nature 1983, 302 (5908), 540–543.

(56) Islam, M. A.; Bhayye, S.; Adeniyi, A. A.; Soliman, M. E. S.; Pillay, T. S. Diabetes Mellitus Caused by Mutations in Human Insulin: Analysis of Impaired Receptor Binding of Insulins Wakayama, Los Angeles and Chicago Using Pharmacoinformatics. Journal of Biomolecular Structure and Dynamics 2017, 35 (4), 724–737.

(57) Hirsch, I. B. Insulin Analogues. New England Journal of Medicine 2005, 352 (2), 174–183.

(58) Drejer, K. The Bioactivity of Insulin Analogues from in Vitro Receptor Binding to in Vivo Glucose Uptake. Diabetes/Metabolism Reviews 1992, 8 (3), 259–285.

(59) Sommerfeld, M. R.; Müller, G.; Tschank, G.; Seipke, G.; Habermann, P.; Kurrle, R.; Tennagels, N. In Vitro Metabolic and Mitogenic Signaling of Insulin Glargine and Its Metabolites. PLOS ONE 2010, 5 (3), e9540.

(60) Gammeltoft, S.; Hansen, B. F.; Dideriksen, L.; Lindholm, A.; Schäffer, L.; Trüb, T.; Dayan, A.; Kurtzhals, P. Insulin Aspart: A Novel Rapid-Acting Human Insulin Analogue. Expert Opinion on Investigational Drugs 1999, 8 (9), 1431–1442.

(61) Holleman, F.; Hoekstra, J. B. L. Insulin Lispro. New England Journal of Medicine 1997, 337 (3), 176–183.

(62) Kurtzhals, Peter, Lauge Schäffer, Anders Sørensen, Claus Kristensen, Ib Jonassen, Christoph Schmid, and Thomas Trüb. “Correlations of receptor binding and metabolic and mitogenic potencies of insulin analogs designed for clinical use.” Diabetes 49, no. 6 (2000): 999–1005.

(63) Kildegaard, J.; Buckley, S. T.; Nielsen, R. H.; Povlsen, G. K.; Seested, T.; Ribel, U.; Olsen, H. B.; Ludvigsen, S.; Jeppesen, C. B.; Refsgaard, H. H. F.; Bendtsen, K. M.; Kristensen, N. R.; Hostrup, S.; Sturis, J. Elucidating the Mechanism of Absorption of Fast-Acting Insulin Aspart: The Role of Niacinamide. Pharm Res 2019, 36 (3), 49.

(64) A. Zib, I.; Raskin, P. Novel Insulin Analogues and Its Mitogenic Potential. Diabetes, Obesity and Metabolism 2006, 8 (6), 611–620.

(65) A. Glendorf, T.; Knudsen, L.; Stidsen, C. E.; Hansen, B. F.; Hegelund, A. C.; Sørensen, R.; Nishimura, E.; Kjeldsen, T. Systematic Evaluation of the Metabolic to Mitogenic Potency Ratio for B10-Substituted Insulin Analogues. PLOS ONE 2012, 7 (2), e29198.

(66) Brange, J.; Ribel, U.; Hansen, J. F.; Dodson, G.; Hansen, M. T.; Havelund, S.; Melberg, S. G.; Norris, F.; Norris, K.; Snel, L.; Sørensen, A. R.; Voigt, H. O. Monomeric Insulins Obtained by Protein Engineering and Their Medical Implications. Nature 1988, 333 (6174), 679–682.

(67) Mukherjee, S.; Mondal, S.; Deshmukh, A. A.; Gopal, B.; Bagchi, B. What Gives an Insulin Hexamer Its Unique Shape and Stability? Role of Ten Confined Water Molecules. J. Phys. Chem. B 2018, 122 (5), 1631–1637.

(68) Schwartz, Gerald P., G. Thompson Burke, and Panayotis G. Katsoyannis. “A superactive insulin:[B10-aspartic acid] insulin (human).” Proceedings of the National Academy of Sciences 84, no. 18 (1987): 6408–6411.

(69) Hansen, B. F.; Kurtzhals, P.; Jensen, A. B.; Dejgaard, A.; Russell-Jones, D. Insulin X10 Revisited: A Super-Mitogenic Insulin Analogue. Diabetologia 2011, 54 (9), 2226–2231.

(70) Glendorf, T.; Sørensen, A. R.; Nishimura, E.; Pettersson, I.; Kjeldsen, T. Importance of the Solvent-Exposed Residues of the Insulin B Chain α-Helix for Receptor Binding. Biochemistry 2008, 47 (16), 4743–4751.

(71) Slieker, L. J.; Brooke, G. S.; DiMarchi, R. D.; Flora, D. B.; Green, L. K.; Hoffmann, J. A.; Long, H. B.; Fan, L.; Shields, J. E.; Sundell, K. L.; Surface, P. L.; Chance, R. E. Modifications in the B10 and B26–30 Regions of the B Chain of Human Insulin Alter Affinity for the Human IGF-I Receptor More than for the Insulin Receptor. Diabetologia 1997, 40 (2), S54–S61.

(72) Glidden, M. D.; Yang, Y.; Smith, N. A.; Phillips, N. B.; Carr, K.; Wickramasinghe, N. P.; Ismail-Beigi, F.; Lawrence, M. C.; Smith, B. J.; Weiss, M. A. Solution Structure of an Ultra-Stable Single-Chain Insulin Analog Connects Protein Dynamics to a Novel Mechanism of Receptor Binding. Journal of Biological Chemistry 2018, 293 (1), 69–88.

(73) Liu, M.; Weiss, M. A.; Arunagiri, A.; Yong, J.; Rege, N.; Sun, J.; Haataja, L.; Kaufman, R. J.; Arvan, P. Biosynthesis, Structure, and Folding of the Insulin Precursor Protein. Diabetes, Obesity and Metabolism 2018, 20 (S2), 28–50.

(74) Rege, N. K.; Wickramasinghe, N. P.; Tustan, A. N.; Phillips, N. F. B.; Yee, V. C.; Ismail-Beigi, F.; Weiss, M. A. Structure-Based Stabilization of Insulin as a Therapeutic Protein Assembly via Enhanced Aromatic–Aromatic Interactions. Journal of Biological Chemistry 2018, 293 (28), 10895–10910.

(75) Akbarian, M.; Yousefi, R.; Moosavi-Movahedi, A. A.; Ahmad, A.; Uversky, V. N. Modulating Insulin Fibrillation Using Engineered B-Chains with Mutated C-Termini. Biophysical Journal 2019, 117 (9), 1626–1641.

(76) Nakagawa, S. H.; Tager, H. S. Role of the Phenylalanine B25 Side Chain in Directing Insulin Interaction with Its Receptor. Steric and Conformational Effects. Journal of Biological Chemistry 1986, 261 (16), 7332–7341.

(77) Fischer, W. H.; Saunders, D.; Brandenburg, D.; Wollmer, A.; Zahn, H. A Shortened Insulin with Full in Vitro Potency. 1985, 366 (1), 521–526.

(78) Casaretto, M.; Spoden, M.; Diaconescu, C.; Gattner, H.-G.; Zahn, H.; Brandenburg, D.; Wollmer, A. Shortened Insulin with Enhanced in Vitro Potency. 1987, 368 (1), 709–716. 10.1515/bchm3.1987.368.1.709.

(79) Sims, E. K.; Carr, A. L. J.; Oram, R. A.; DiMeglio, L. A.; Evans-Molina, C. 100 Years of Insulin: Celebrating the Past, Present and Future of Diabetes Therapy. Nat Med 2021, 27 (7), 1154–1164.

(80) Menting, J. G.; Lawrence, C. F.; Kong, G. K.-W.; Margetts, M. B.; Ward, C. W.; Lawrence, M. C. Structural Congruency of Ligand Binding to the Insulin and Insulin/Type 1 Insulin-like Growth Factor Hybrid Receptors. Structure 2015, 23 (7), 1271–1282.

(81) Li, J.; Choi, E.; Yu, H.; Bai, X. Structural Basis of the Activation of Type 1 Insulin-like Growth Factor Receptor. Nat Commun 2019, 10 (1), 4567.

(82) Kitagawa, K.; Ogawa, H.; Burke, G. T.; Chanley, J. D.; Katsoyannis, P. G. Interaction between the A2 and A19 Amino Acid Residues Is of Critical Importance for High Biological Activity in Insulin: [19-Leucine-A]Insulin. Biochemistry 1984, 23 (19), 4444–4448.

(83) Kristensen, C.; Kjeldsen, T.; Wiberg, F. C.; Schäffer, L.; Hach, M.; Havelund, S.; Bass, J.; Steiner, D. F.; Andersen, A. S. Alanine Scanning Mutagenesis of Insulin *. Journal of Biological Chemistry 1997, 272 (20), 12978–12983.

(84) Kertisová, A.; Žáková, L.; Macháčková, K.; Marek, A.; Šácha, P.; Pompach, P.; Jiráček, J.; Selicharová, I. Insulin Receptor Arg717 and IGF-1 Receptor Arg704 Play a Key Role in Ligand Binding and in Receptor Activation. Open Biol 13 (11), 230142.

(85) Shoelson, S.; Fickova, M.; Haneda, M.; Nahum, A.; Musso, G.; Kaiser, E. T.; Rubenstein, A. H.; Tager, H. Identification of a Mutant Human Insulin Predicted to Contain a Serine-for-Phenylalanine Substitution. Proceedings of the National Academy of Sciences 1983, 80 (24), 7390–7394.

(86) Kobayashi, M.; Haneda, M.; Maegawa, H.; Watanabe, N.; Takada, Y.; Shigeta, Y.; Inouye, K. Receptor Binding and Biological Activity of [SerB24]-Insulin, an Abnormal Mutant Insulin. Biochemical and Biophysical Research Communications 1984, 119 (1), 49–57.

(87) Heinemann, L.; Starke, A. A. R.; Heding, L.; Jensen, I.; Berger, M. Action Profiles of Fast Onset Insulin Analogues. Diabetologia 1990, 33 (6), 384–386.

(88) Bolli, G. B., R. D. Di Marchi, G. D. Park, S. Pramming, and Veikko A. Koivisto. “Insulin analogues and their potential in the management of diabetes mellitus.” Diabetologia 42, no. 10 (1999): 1151.

(89) Bolli, G. B.; Owens, D. R. Insulin Glargine. The Lancet 2000, 356 (9228), 443–445.

(90) Falck Hansen, B.; Danielsen, G. M.; Drejer, K.; Sørensen, A. R.; Wiberg, F. C.; Klein, H. H.; Lundemose, A. G. Sustained Signalling from the Insulin Receptor after Stimulation with Insulin Analogues Exhibiting Increased Mitogenic Potency. Biochemical Journal 1996, 315 (1), 271–279.

(91) Spoden, M.; Gattner, H.-G.; Zahn, H.; Brandenburg, D. Structure-Function Relationships of Des-(B26-B30)-Insulin. International Journal of Peptide and Protein Research 1995, 46 (3–4), 221–227.

(92) Kurapkat, G.; Siedentop, M.; Gattner, H.-G.; Hagelstein, M.; Brandenburg, D.; Grötzinger, J.; Wollmer, A. The Solution Structure of a Superpotent B-Chain-Shortened Single-Replacement Insulin Analogue. Protein Science 1999, 8 (3), 499–508.

(93) Cavasotto, C. N.; Phatak, S. S. Homology Modeling in Drug Discovery: Current Trends and Applications. Drug Discovery Today 2009, 14 (13), 676–683.

(94) Šali, A.; Blundell, T. L. Comparative Protein Modelling by Satisfaction of Spatial Restraints. Journal of Molecular Biology 1993, 234 (3), 779–815.

(95) Gómez, J.; Freire, E. Thermodynamic Mapping of the Inhibitor Site of the Aspartic Protease Endothiapepsin. Journal of Molecular Biology 1995, 252 (3), 337–350.

(96) Li, H.; Robertson, A. D.; Jensen, J. H. Very Fast Empirical Prediction and Rationalization of Protein PKa Values. Proteins: Structure, Function, and Bioinformatics 2005, 61 (4), 704–721. 10.1002/prot.20660.

(97) Sham, Yuk Yin, Zhen Tao Chu, and Arieh Warshel. “Consistent calculations of p K a’s of ionizable residues in proteins: semi-microscopic and microscopic approaches.” The Journal of Physical Chemistry B 101, no. 22 (1997): 4458–4472.

(98) Jorgensen, W. L.; Chandrasekhar, J.; Madura, J. D.; Impey, R. W.; Klein, M. L. Comparison of Simple Potential Functions for Simulating Liquid Water. J. Chem. Phys. 1983, 79 (2), 926–935.

(99) Maier, J. A.; Martinez, C.; Kasavajhala, K.; Wickstrom, L.; Hauser, K. E.; Simmerling, C. Ff14SB: Improving the Accuracy of Protein Side Chain and Backbone Parameters from Ff99SB. J Chem Theory Comput 2015, 11 (8), 3696–3713..

(100) Case, David A., Thomas E. Cheatham III, Tom Darden, Holger Gohlke, Ray Luo, Kenneth M. Merz Jr, Alexey Onufriev, Carlos Simmerling, Bing Wang, and Robert J. Woods. “The Amber biomolecular simulation programs.” Journal of computational chemistry 26, no. 16 (2005): 1668–1688.

(101) Kräutler, V.; van Gunsteren, W. F.; Hünenberger, P. H. A Fast SHAKE Algorithm to Solve Distance Constraint Equations for Small Molecules in Molecular Dynamics Simulations. Journal of Computational Chemistry 2001, 22 (5), 501–508.

(102) Essmann, U.; Perera, L.; Berkowitz, M. L.; Darden, T.; Lee, H.; Pedersen, L. G. A Smooth Particle Mesh Ewald Method. J. Chem. Phys. 1995, 103 (19), 8577–8593.

(103) Berendsen, H. J. C.; Postma, J. P. M.; van Gunsteren, W. F.; DiNola, A.; Haak, J. R. Molecular Dynamics with Coupling to an External Bath. J. Chem. Phys. 1984, 81 (8), 3684– 3690.

(104) Ryckaert, J.-P.; Ciccotti, G.; Berendsen, H. J. C. Numerical Integration of the Cartesian Equations of Motion of a System with Constraints: Molecular Dynamics of n-Alkanes. Journal of Computational Physics 1977, 23 (3), 327–341.

(105) Bauer, P.; Barrozo, A.; Purg, M.; Amrein, B. A.; Esguerra, M.; Wilson, P. B.; Major, D. T.; Åqvist, J.; Kamerlin, S. C. L. Q6: A Comprehensive Toolkit for Empirical Valence Bond and Related Free Energy Calculations. SoftwareX 2018, 7, 388–395.

(106) Singh, N.; Warshel, A. Absolute Binding Free Energy Calculations: On the Accuracy of Computational Scoring of Protein-Ligand Interactions. Proteins 2010, 78 (7), 1705–1723.

(107) Marelius, J.; Kolmodin, K.; Feierberg, I.; Åqvist, J. Q: A Molecular Dynamics Program for Free Energy Calculations and Empirical Valence Bond Simulations in Biomolecular Systems11Color Plate 1, Color Plate 2 for This Article Are on Page 261. Journal of Molecular Graphics and Modelling 1998, 16 (4), 213–225.

(108) Lee, F. S.; Warshel, A. A Local Reaction Field Method for Fast Evaluation of Long-range Electrostatic Interactions in Molecular Simulations. J. Chem. Phys. 1992, 97 (5), 3100–3107.

(109) Kollman, P. Free energy calculations: Applications to chemical and biochemical phenomena. ACS Publications.

