## Supplementary file for "Mapping Structural Drivers of Insulin Analogs Using Molecular Dynamics and Free Energy Calculations at Insulin Receptor"

Table S1**:** A summary of currently available XRD and Cryo-EM structures for IR receptors in complex with insulin or as apo receptors.

| **Receptor** | | **Method** | **PDB IDs** | **#SS** | **Year** | **Binding Site** | **Ref.** |
|---|---|---|---|---|---|---|---|
| IR | High affinity insulin analogs | XRD | 3W12<br>3W13<br>5KQV(3W14) | 3<br>3 | 2013<br>2013 | HA<br>HA | ([18])<br>([18]) |
| IR | Insulin (wild type) | XRD | 4OGA | 3 | 2014 | HA | ([28]) |
| IR-apo | - | XRD | 4ZXB | - | 2016 | - | ([29]) |
| IR | Insulin (wild type) | EM | 6CE9 | 1 | 2018 | HA | ([25]) |
| | | | 6CEB | 1 | 2018 | HA | ([25]) |
| | | | 6PXV | 3 | 2019 | Both | ([26]) |
| | | | 6PXW | 3 | 2019 | Both | ([26]) |
| | | | 6SOF | 1/3 | 2020 | HA/LA | ([2]) |
| | | | 7QID | 3 | 2020 | HA/LA | ([2]) |
| | | | 7PG4 | 2 | 2022 | HA/LA | ([27]) |

Note**:** #SS: Number of disulfide bridges in Ligand. BS: binding site type. HA: High affinity and LA: Low affinity

Table S2**:** Binding enthalpies calculated from MM-GBSA calculation. Mean, SEM and Sd. are calculated from the ΔH calculated from ligand molecule bound at two HA sites, s1 & s1' from two simulations with different initial velocities.

| Complex | $\Delta H_{bind}$ (kcal/mol) (MM-GBSA) | | SASA(Å$^2$) (LCPO, surface) | |
|---|---|---|---|---|
| | Mean | SEM | Mean | Std. Dev. |
| 6CE9: WT-Ins | -66.29 | 8.28 | 618.70 | 74.96 |

Table S3: FEP scheme summary sheet

| Insulin Analog | FEP transformation | Intermediate Exists | Intermediate State | Total No. of steps involved in electrostatic transformation | Total No. of steps involved Softcore potentials | Total No. of steps involved in Van der Waals transformation |
|---|---|---|---|---|---|---|
| Wakayma | V-L | Yes | Alanine | 74 | 44 | 252 |
| Los Angles | F-S | Yes | Alanine | 94 | 44 | 252 |
| Chicago | F-L | Yes | Cγ | 44 | 44 | 168 |
| B10Q | H-Q | Yes | Cγ | 144 | 44 | 168 |
| B10A | H-A | No | NA | 72 | 22 | 84 |
| B10V | H-V | Yes | Alanine | 144 | 44 | 210 |
| B10D | H-D | No | NA | 144 | 44 | 168 |
| A21G | N-G | No | NA | 72 | 22 | 168 |
| Des-B25-F | H-F | Yes | Cγ | 144 | 44 | 210 |
| Des-B25-Y | Y-H | Yes | Cγ | 144 | 42 | 168 |
| Des-B25-W | W-H | No | NA | 72 | 42 | 168 |

Table S4: Calculated relative free energies ($\Delta\Delta G_{bind}$) at IR for all the insulin analogs considered in this study along with the range of experimental values available for each analog. The experimental spread of $\Delta\Delta G_{bind}$ has been calculated using the inner means and a mean of means has been considered in order to compare with the calculated $\Delta\Delta G_{bind}$. The experimental and calculated $\Delta\Delta G_{bind}$ have both been reported in relation to WT-Ins or equivalent in the case of truncated systems.

| Transformation | State-1 | State-2 | Experimental ΔΔG (mean)(kcal/mol) | FEP ΔΔG (kcal/mol) | Trend | Error (kcal/mol) (exp-sim) |
|---|---|---|---|---|---|---|
| Wakayama | Valine | Alanine | 3.1 | 0.61 | Yes (orange) | 2.5 |
| Los Angles | Phenyl Alanine | Serine | 1.5 | -0.56 | No (red) | 2.1 |
| Chicago | Phenyl Alanine | Leucine | 2.3 | 0.79 | Yes (orange) | 1.5 |
| Lispro | PKT | KPT | 0.12 | -0.43 | Yes (dark green) | 0.55 |
| Aspart | DKT | PKT | -0.0060 | -0.3 | Yes (dark green) | 0.29 |
| A21Gly | Aspargine | Glycine | 0.079 | 0.71 | Yes (dark green) | -0.63 |
| B10Asp | Histidine | Aspartic acid | -0.45 | -1.64 | Yes (orange) | 1.16 |
| B10V | Histidine | Valine | 0.24 | 0.28 | Yes (dark green) | -0.04 |
| B10A | Histidine | Alanine | 0.18 | 1.05 | Yes (dark green) | -0.85 |
| B10Q | Histidine | Glutamine | -0.01 | -0.5 | Yes (dark green) | 0.49 |
| B25H* | Phenyl Alanine | Histidine | -0.7 | -1.53 | Yes (dark green) | +0.8 |
| B25Y* | Phenyl Alanine | Tyrosine | -0.57 | -1.9 | Yes (orange) | +1.4 |
| B25W* | Phenyl Alanine | Tryphtophan | 0.38 | -2.2 | No (red) | 2.58 |

*: Truncated Insulins (desB26-B30)

Table S5: Ensemble averaged solvent interaction networks for B10Asp & B10His in IR from MD trajectories.

| | Protein | | Water | |
|---|---|---|---|---|
| | IR | | IR | |
| | B10Asp | B10His | B10Asp | B10His |
| Zero order contacts | 2.24 (P0) | 0.06 (P0) | 4.38 (W0) | 2.84 (W0) |
| First order contacts | 2.68 (P1) | 0.76 (P1) | 13.08 (W1) | 9.16 (W1) |
| Second order contacts | 28.16 (P2) | 8.26 (P2) | 40.10 (W2) | 15.98 (W2) |

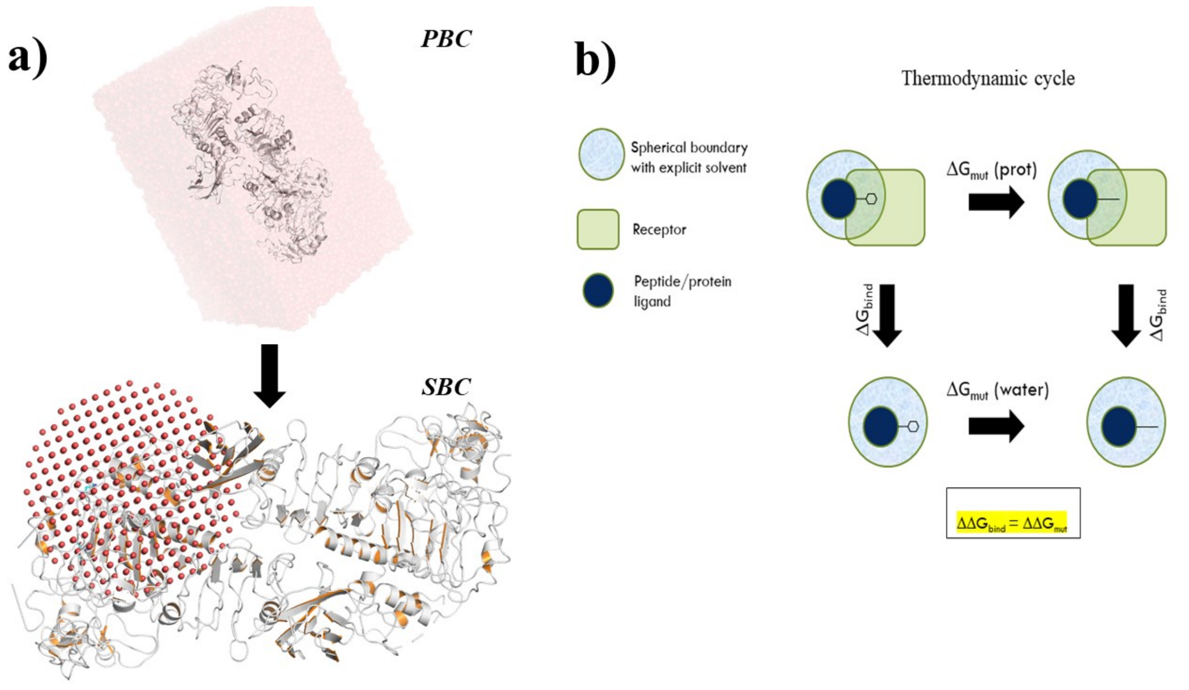


**Figure S1:** a) depicts periodic boundary conditions (PBC) used for MD simulations and spherical boundary conditions (SBC) used for MD-FEP calculations. The top panel shows water molecules around the protein (cartoon) as surface representation in the PBC simulations. The bottom panel shows SBC used for FEP calculations in this study. All atoms within the sphere are fully flexible whereas those outside the sphere are restrained to their initial coordinates. The water molecules are shown as red spheres with receptors shown as cartoons. b) Schematic representation of thermodynamic cycle used for FEP calculations. The blue circle represents the sphere, green rounded rectangle represents receptor and filled dark blue circle represents the peptide ligand. The same mutations are performed with the peptide bound to the receptor and with free peptide in water. The relative $\Delta\Delta G_{bind}$ is calculated by taking advantage of the thermodynamic cycle as shown.
